# Developmental convergence and divergence in human stem cell models of autism spectrum disorder

**DOI:** 10.1101/2024.04.01.587492

**Authors:** Aaron Gordon, Se-Jin Yoon, Lucy K Bicks, Jaqueline M Martin, Greta Pintacuda, Stephanie Arteaga, Brie Wamsley, Qiuyu Guo, Lubayna Elahi, Ricardo E. Dolmetsch, Jonathan A Bernstein, Ruth O’Hara, Joachim F Hallmayer, Kasper Lage, Sergiu P Pasca, Daniel H Geschwind

**Affiliations:** Program in Neurogenetics, Department of Neurology, David Geffen School of Medicine, University of California Los Angeles, Los Angeles, CA, USA; Department of Psychiatry and Behavioral Sciences, Stanford University, Stanford, CA, USA; Stanford Brain Organogenesis Program, Wu Tsai Neurosciences Institute and Bio-X, Stanford University, Stanford, CA, USA; Department of Human Genetics, David Geffen School of Medicine, University of California Los Angeles, Los Angeles, CA, USA; Program in Neurobehavioral Genetics, Semel Institute, David Geffen School of Medicine, University of California Los Angeles, Los Angeles, CA, USA; Center for Autism Research and Treatment, Semel Institute, David Geffen School of Medicine, University of California Los Angeles, Los Angeles, CA, USA; Tempero Bio, Boston, Massachusetts, USA; Department of Pediatrics, Stanford University, Stanford, CA, USA; Stanley center for Psychiatric Research at the Broad Institute

## Abstract

Two decades of genetic studies in autism spectrum disorder (ASD) have identified over a hundred genes harboring rare risk mutations. Despite this substantial heterogeneity, transcriptomic and epigenetic analyses have identified convergent patterns of dysregulation across ASD post-mortem brain tissue. To identify shared and distinct mutational mechanisms, we assembled the largest hiPS cell patient cohort to date, consisting of 70 hiPS cell lines after stringent quality control representing 8 ASD-associated mutations, idiopathic ASD, and 20 lines from non-affected controls. We used these hiPS lines to generate human cortical organoids (hCO), profiling by RNAseq at four distinct timepoints up to 100 days of *in vitro* differentiation. Early timepoints harbored the largest mutation-specific changes, but different genetic forms converged on shared transcriptional changes as development progressed. We identified a shared RNA and protein interaction network, which was enriched in ASD risk genes and predicted to drive the observed down-stream changes in gene expression. CRISPR-Cas9 screening of these candidate transcriptional regulators in induced human neural progenitors validated their downstream molecular convergent effects. These data illustrate how genetic risk can propagate via transcriptional regulation to impact convergently dysregulated pathways, providing new insight into the convergent impact of ASD genetic risk on human neurodevelopment.

## Introduction

Autism spectrum disorder (ASD) is a common neurodevelopmental disorder, with a childhood prevalence of close to 2% (Zeidan et al., 2022). The last decade of genetic studies has yielded hundreds of risk genes, consistent with extraordinary etiological heterogeneity, which poses enormous challenges for investigating the mechanisms underlying ASD (de la Torre-Ubieta et al., 2016; Huguet et al., 2013; Satterstrom et al., 2020; Willsey et al., 2022). Over 100 high confidence genetic mutations have been associated with ASD in genetic studies (De Rubeis et al., 2014; Feliciano et al., 2019; Iossifov et al., 2014; O’Roak et al., 2012; Ruzzo et al., 2019a; Sanders et al., 2015; Satterstrom et al., 2020; Sebat et al., 2007; Werling et al., 2018; Yuen et al., 2017). These genes, which carry rare, often *de novo*, single nucleotide variations (SNVs), or copy number variants (CNVs) with moderate to large effect sizes, are observed in 10%-20% of ASD cases (Geschwind and State, 2015; Havdahl et al., 2021). Common variation is predicted to explain another 50% of the genetic risk for ASD (Gaugler et al., 2014; Grove et al., 2019; Weiner et al., 2017) however to date only five common loci have been significantly associated with ASD (Grove et al., 2019). Based on this complex genetic architecture, an important component of ASD risk can be construed as a collection of distinct, rare disorders with highly overlapping clinical features.

Despite this heterogeneity in the genetic etiology of ASD, post-mortem transcriptome analysis has revealed consistent differences among a majority of subjects with idiopathic ASD, as well as subjects with a specific syndromic form of ASD, (dup)15q11-13 (Gandal et al., 2022; Parikshak et al., 2016; Velmeshev et al., 2019). However, the mechanism by which multiple distinct mutations can lead to convergent molecular pathology, and whether molecular convergence occurs across other rare forms of ASD, remains unknown. Understanding this progression is complicated by the fact that many ASD-associated mutations are in genes whose expression peaks during fetal development, while gene expression studies are conducted after this critical developmental window has ended (Gandal et al., 2018; Gandal et al., 2022; Gupta et al., 2014; Parikshak et al., 2013; Willsey et al., 2013). Further, several lines of evidence, including genetic (Ben-David and Shifman, 2013), genomic (Walker et al., 2019; Won et al., 2019), neuroimaging (Hazlett et al., 2017), and neuropathology (Chen et al., 2015), indicate that early neurodevelopment plays an essential role in the development of ASD and other neuropsychiatric disorders (Heckers et al., 2015).

The advent of stem cell-based *in vitro* systems enables modeling potential disruptions of human brain development in neurodevelopmental disorders (Amin and Paşca, 2018; Gordon and Geschwind, 2020; Pașca et al., 2022; Seah et al., 2023). In particular, three-dimensional (3D) regionalized neural organoids were found to model several aspects of *in vivo* human neurodevelopment (Amiri et al., 2018; Gordon et al., 2021; Paşca et al., 2015; Velasco et al., 2019). Crucially, this allows us to emulate previously inaccessible early neurodevelopmental stages across time within an individual, opening a new field of assessing the genetic basis of altered neurodevelopmental trajectories instead of relying on traditional cross-sectional approaches (Arlotta and Paşca, 2019; Pașca, 2018; Seah et al., 2023). Most studies have investigated relatively small numbers of lines from patients with idiopathic ASD (Jourdon et al., 2023), or have focused on individual mutations, which has demonstrated the utility of hiPS cell-based systems to study the impact of ASD genetic risk on neurodevelopmental processes (Birey et al., 2022; Khan et al., 2020; Lancaster et al., 2013; Paulsen et al., 2022). Additionally, recent work has highlighted the power of studying the impact of multiple genes in parallel, identifying additional evidence for molecular convergence in ASD (Jin et al., 2020; Li et al., 2023; Meng et al., 2023).

Here, we profile the largest hiPS cell patient cohort to date, starting from 96 hiPS cell lines in total from individuals with 8 different mutations associated with ASD, and from 11 individuals with idiopathic ASD and 30 lines derived from 25 matched controls, which after stringent QC yielded 70 lines for downstream analysis. From each of these lines, we derived neural organoids using a guided differentiation approach that we developed to resemble the cerebral cortex, known as human cortical organoids (hCOs) (Paşca et al., 2015; Yoon et al., 2019). We measure their transcriptional profiles, which represent reliable quantitative phenotypes, at multiple stages of development up to 100 days of differentiation *in vitro*. We demonstrate that these cases of ASD with distinct genetic risk, which we refer to as “different forms” of ASD, while showing distinct profiles, also reliably cluster based on changes in gene expression. Using orthogonal analysis approaches including differential expression, independent component analysis and co-expression network analysis, we find evidence for convergence onto changes in early neuronal differentiation. We identify and characterize a downregulated, chromatin and transcriptional network which contains several ASD risk genes, including members of the SWI/SNF complex. This network is predicted to be upstream and drive the observed changes in downstream gene expression associated with these ASD risk mutations. Lastly, we use CRISPRi to validate the effects of multiple putative network drivers on downstream gene expression, which includes down-regulation of multiple ASD risk genes and key pathways in neural development.

## Results

### Generation of hCOs from hiPS cell lines

We reprogrammed somatic cells (fibroblasts or peripheral blood mononuclear cells (PBMCs); Methods) to generate human induced pluripotent stem cells (hiPS cells) from a cohort of individuals with 8 different mutations associated with ASD: (1) 22q11.2 deletion, a 3 Mb deletion which leads to a constellation of variably present symptoms including heart defects, palate and other craniofacial features, intellectual disability, ASD (∼20%) and psychosis (∼25%) (McDonald-McGinn et al., 2015; Swillen and McDonald-McGinn, 2015)(n = 18 (Khan et al., 2020)); (2) 22q13.3 deletion, a deletion spanning 130 kb to 9 Mb which includes *SHANK3, ACR,* and *RABL2* genes among others and presents with developmental delay, hypotonia, delayed speech, and impaired social interactions (Phelan and McDermid, 2011)(n = 4); (3) 15q13.3 deletion, a 1.5 Mb deletion spanning the six genes *MTMR15, MTMR10, TRPM1, KLF13, OTUD7A, and CHRNA7* that typically presents with intellectual disability and epilepsy (Sharp et al., 2008) (n = 3); (4) 16p11.2 deletion, a ∼600 kb deletion presenting with intellectual disability, motor impairments, speech and communication deficits, and ASD (Chung et al., 2021) (n = 4); (5) 16p11.2 duplication, which shares intellectual disability and ASD phenotypes with the reciprocal deletion (Steinman et al., 2016) (n = 4); (6) Timothy syndrome, which is characterized by variants within the *CACNA1C* gene and presents with syndactyly, prolonged QT interval, ASD and intellectual disability (Splawski et al., 2004) (Bauer et al., 2021) (n = 2 (Birey et al., 2022; Paşca et al., 2011; Yazawa et al., 2011)); (7) PCDH19-related disorder, which is associated with epilepsy and can also include intellectual disability and ASD (Gursoy et al., 2020; Ryan et al., 1997) (n = 2); and (8) SHANK3 R522W mutation (n = 1), a point mutation in SHANK3 associated with neurodevelopmental risk (Gauthier et al., 2010); other pathogenic SHANK3 mutations share behavioral phenotypes with the larger 22q13.3 deletion (De Rubeis et al., 2018; Schön et al., 2023; Yazawa et al., 2011), as well as individuals with idiopathic ASD (n = 11) and unaffected individuals (n = 25) (Figure 1a and Supplementary Table 1) for a total of 74 individuals. For several individuals we used multiple hiPS cell lines, which we used to assess reproducibility, amounting to a total of 96 hiPS cell lines (Figure 1a-b and Extended Data Figure 1).

**Figure 1.**
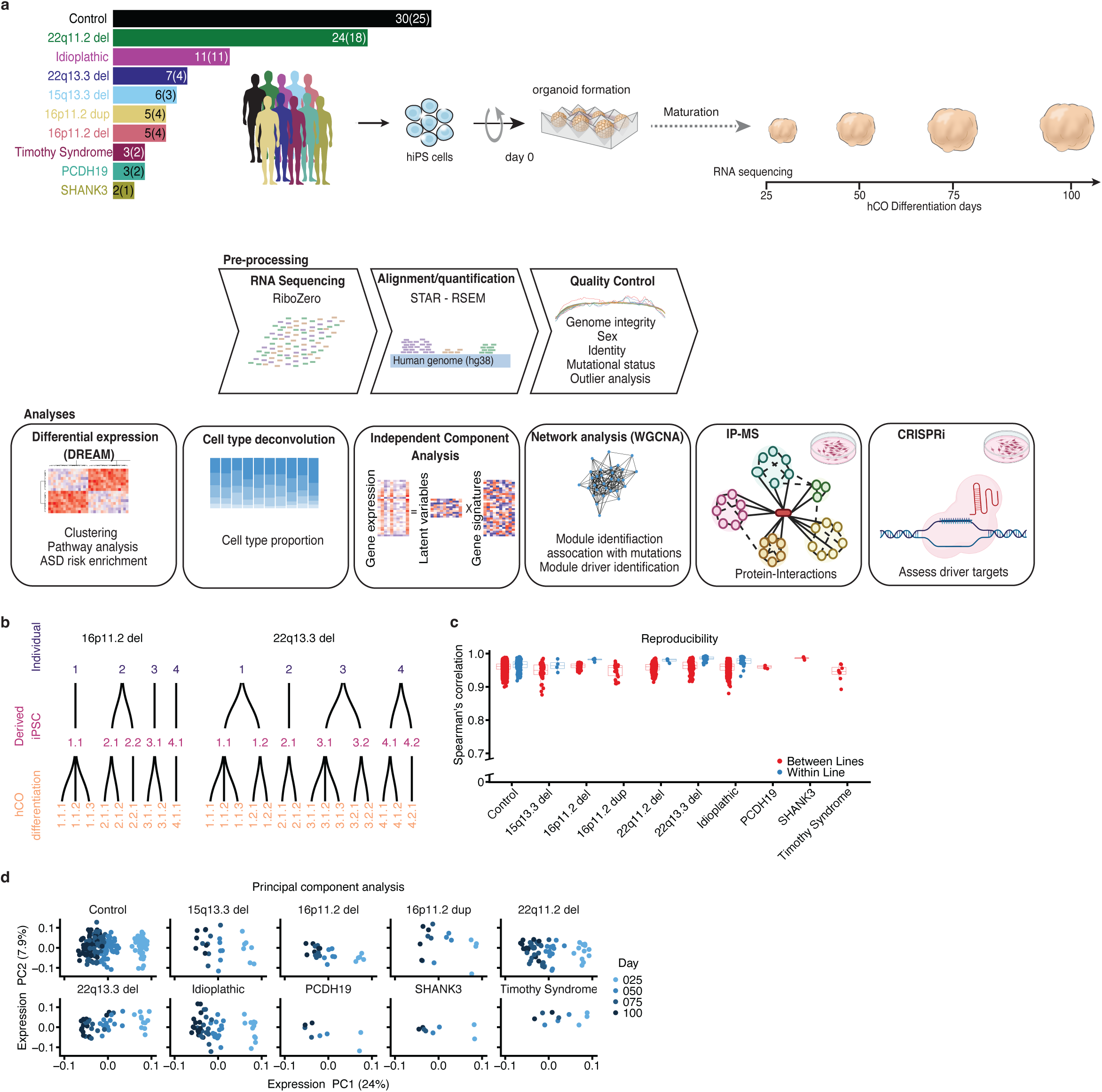
Experimental workflow and validation. (a) Schematic workflow going from hiPS cells to cortical organoids to sequencing data. Number of hiPS cells lines and individuals (in brackets) for each form of ASD and controls is indicated. (b) Schematic representation of hCO differentiations derived from each hiPS cells line for two forms of ASD (16p11.2 deletion and 22q13.3 deletion). The other forms of ASD can be found in Extended data Figure 1. (c) Spearman’s correlation of gene expression between samples from the same timepoint and form of ASD that were derived either from different individuals (red) or from the same individual (blue). Boxplots show: center – median, lower hinge – 25% quantile, upper hinge – 75% quantile, lower whisker – smallest observation greater than or equal to lower hinge –1.5× interquartile range, upper whisker – largest observation less than or equal to upper hinge +1.5× interquartile range. (d) Scatter plot of the first two gene expression principal components (PC) from each form of ASD. Color represents the differentiation day. PC1 explains 24% of the variation and is highly associated with differentiation day. PC2 explains 7.9% of the variation.

We differentiated hiPS cell lines into hCO for 100 days using a scalable and reproducible guided differentiation method (Yoon et al., 2019), which yields hCO that contain developing radial glia and ultimately give rise to functional glutamatergic neurons in a process that closely resemble the developing cerebral cortex (Gordon et al., 2021; Yoon et al., 2019). In total, we performed 172 unique hCO differentiations of the 96 lines, so as to have replicates for half (49%) of the lines (control n = 75, idiopathic ASD n = 21, 22q11.2 deletion n = 31, 22q13.3 deletion n = 15, 15q13.3 deletion n = 7, 16p11.2 deletion n = 9, 16p11.2 duplication n = 5, Timothy syndrome n = 3, PCDH19 mutation n = 4, and SHANK3 mutation n = 2). We performed whole genome sequencing to verify genome integrity and removed any lines with large CNVs, viral integration, or where the mutation of interest was absent, which yielded 70 verified lines from 55 individuals passing QC for downstream functional genomic analyses (Methods; Extended Data Figures 1,2,3 and Supplementary Table 2).

We next analyzed the transcriptomes of pooled hCOs at day 25, 50, 75 and 100 of hCO differentiation (Figure 1a-b; Extended Data Figure 1). In these cultures, day 25 corresponds to an early period in cortical development (Gordon et al., 2021; Paşca et al., 2015), consisting primarily of cycling progenitors and radial glia (Kanton et al., 2019; Paşca et al., 2015). As differentiation progresses, the cell composition evolves to consist of intermediate progenitors, deep layer post-mitotic neurons and astroglial lineage cells (Khan et al., 2020; Paşca et al., 2015). After stringent RNA-sequencing quality control and outlier analysis to remove low-quality samples (Figure 1a; Extended Data Figure 1-3; Methods), 464 samples from 55 individuals (70 lines) were used for downstream analysis (Supplementary Table 1).

Gene expression provides a highly reliable, quantitative, genome-wide metric of the state of the cells or tissue (2020; Ferraro et al., 2020), thus, we first evaluated gene expression reproducibility across samples (Methods), finding high reproducibility both between (overall mean Spearman correlation 0.96, range 0.88–0.99) and within (overall mean Spearman correlation 0.97, range 0.92–0.98) individuals, similar to published metrics (Gordon et al., 2021; Khan et al., 2020; Yoon et al., 2019) (Figure 1c). Next, we evaluated the drivers of variation in the data and found that the largest driver of variation was stage of differentiation (Figure 1d and Extended Data Figure 4), consistent with our previous observations (Gordon et al., 2021). Gene expression within the CNVs was downregulated in the deletion carriers and upregulated in the duplication carriers across the different time points, as per expectations (Extended Data Figure 5). While several of the genes affected by the point mutations showed a trend towards downregulation (PCDH19: logFC=–0.77, FDR=0.45; SHANK3: logFC= –0.03, FDR=1.0; CACNA1C: logFC= –0.22, FDR=0.78), these changes were not statistically significant; this is expected based on what is known about the pathogenic mechanisms of these missense mutations (Gerosa et al., 2019; Han et al., 2019; Leblond et al., 2014) Extended Data Figure 3 and 5).

### Transcriptomic relationships across mutational type and over differentiation time

To determine the relationship of gene expression patterns across different forms of ASD, we performed hierarchical clustering based on the correlation of log_2_ fold changes in gene expression (Supplementary table 3). We observed several reliable, distinct clusters (Methods), with subjects harboring the 16p11.2 deletion clustering with those having the PCDH19-related disorder, 16p11.2 duplication clustering with Timothy syndrome and 22q13.3 and SHANK3 mutation clustering together (Figure 2a). These clusters were significant and were robust to the method of clustering and to the number of genes used (Extended data Figure 6; Methods).

**Figure 2.**
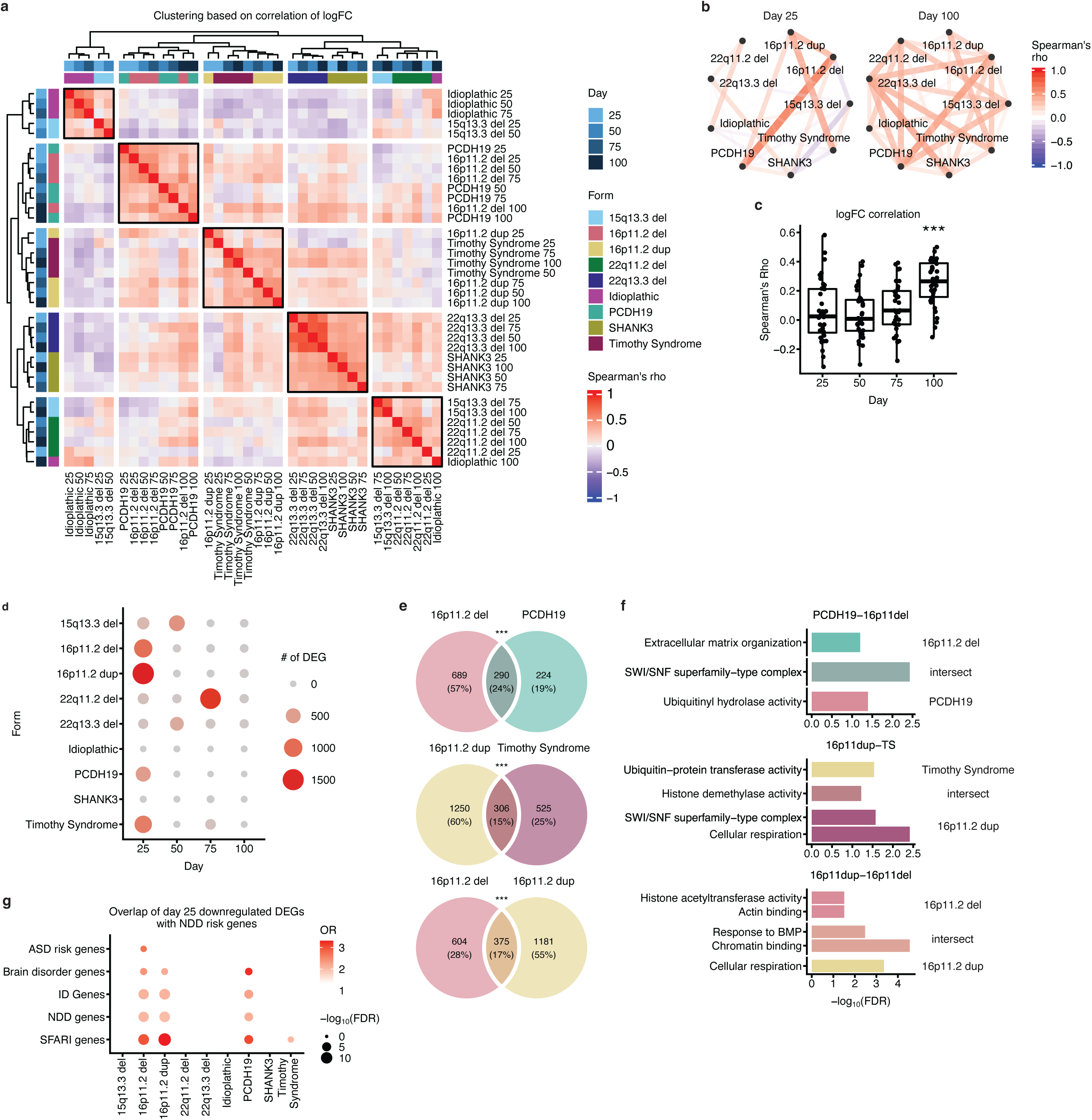
Clustering and overlap of ASD forms points. (a) Heatmap of hierarchical clustering based on correlation of log fold change between the different conditions (ASD form/differentiation day). Top and left most annotation bars (shades of blue) represent the form of ASD. Bottom and left annotation bars (colorful boxes) represent the differentiation day. (b) Network representation of the correlation (Spearman’s rho) between log fold change of the different forms of ASD at day 25 and day 100. Color corresponds to the value of rho and line thickness corresponds to the absolute value of rho. (c) All correlations (Spearman’s rho) between log fold change of the different. ANOVA followed by post-hoc Tukey’s HSD; ANOVA F_3,140_= 10.97, p = 1.6×10^-6^, Tukey HSD: day100-day25p =2.67×10^-5^, day100-day50p =8×10^-6^, day100-day75p =7×10^-4^. Boxplots show: center – median, lower hinge – 25% quantile, upper hinge – 75% quantile, lower whisker – smallest observation greater than or equal to lower hinge –1.5× interquartile range, upper whisker – largest observation less than or equal to upper hinge +1.5× interquartile range. (d) Number of differentially expressed genes in each form of ASD during hCO differentiation. Color and size of circle represent the number of differentially expressed genes. (e) Overlap in the differentially expressed genes between forms of ASD at day 25. Number and percent of total genes (in brackets) are indicated in each section of the Venn diagram. (f) GO terms significantly enriched in unique and intersecting differentially expressed genes. (g) Overlaps between differentially expressed genes at day 25 and risk genes for ASD from either SFARI or from large scale whole exome sequencing (WES) studies (Ruzzo et al., 2019a; Satterstrom et al., 2020) as well as with neurodevelopmental disorders (NDDs) and intellectual disability (ID) risk genes (Leblond et al., 2021). Color represents the OR and the size of the point represented the −log_10_FDR. Only positive significant overlaps (OR > 1 and FDR < 0.05) are shown. *** FDR < 0.001.

Despite the strong effect of differentiation day on transcriptional profiles, most samples from individual genetic events mutations clustered together. One interesting exception was 15q13.3 for which later timepoints (day 75 and day 100) clustered with 22q11.2 deletion, while earlier time points (day 25 and day 50) clustered with idiopathic ASD (Figure 2a). Examining the correlation of log_2_ fold-changes by differentiation stage revealed that the correlations between hCO harboring different genetic events were significantly stronger at day 100 (ANOVA F_3,140_= 11, p=1.6×10^-6^; Tukey HSD: day100-day25 p=2.7×10^-5^, day100-day50 p=8.0×10^-6^, day100-day75 p=7.0×10^-4^; Figure 2b-c) suggesting that the changes in gene expression converge as cortical differentiation progresses *in vitro.* Furthermore, the number of differentially expressed genes (FDR < 0.05) was lowest on day 100 across all forms of ASD (Figure 2d), suggesting that the differences between cases and controls become less pronounced with time, consistent with the idea of buffering and canalization as development progresses (Kadelka et al., 2024) (Debat and David, 2001; Farrell et al., 2018; Takahashi, 2019).

Next, by examining the number of differentially expressed genes at each time point, we found that the largest changes in gene expression were observed at early stages of hCO differentiation with the largest number of differentially expressed genes found at day 25 in four out of the nine mutational classes (16p11.2 deletion – 979 genes, 16p11.2 duplication – 1556 genes, PCDH19 – 514 genes, and Timothy syndrome – 832 genes), or at day 50 in two of the remaining forms (15q13.3 deletion - 616 genes, and 22q13.3 deletion - 373 genes; Figure 2d and Supplementary table 3). Unlike the large effects found in the above mutations, we detected only 2 genes significantly differentially expressed in the idiopathic cohort (*PRRC2C* at day 50 and the lncRNA *RP11-114H21.2* at day 100; FDR<0.05). This is likely due to the heterogeneity of idiopathic ASD as well as a potentially protracted developmental emergence and reflects the need for larger sample size to detect signal.

The four forms of ASD that had the largest changes at day 25 (16p11.2 deletion, 16p11.2 duplication, PCDH19-related disorders, and Timothy syndrome) formed 2 adjacent clusters (Figure 2a), within which differentially expressed genes significantly intersected (16p11.2 deletion and PCDH19 mutation – 290/1203 genes, 16p11.2 duplication and Timothy syndrome – 306/2081 genes, p<10^-16^ for both, hypergeometric test). Significant similarity was also seen between genes differentially expressed in samples harboring deletion and duplication of 16p11.2 (375/2160 genes, p<10^-16^, hypergeometric test) (Figure 2e). The intersecting genes were significantly enriched for chromatin remodeling related terms (*“SWI/SNF superfamily-type complex”, “histone demethylase activity” and “chromatin binding*”) (Figure 2f), which have previously been associated with ASD risk genes (Mossink et al., 2021; Parikshak et al., 2013; Ruzzo et al., 2019a). Remarkably, genes down-regulated at 25 days in these four forms of ASD were enriched for known high confidence ASD risk genes and those more broadly involved in neurodevelopmental disorders (16p11.2 del: SFARI genes OR=3.0, FDR=3.8×10^-9^; 16p11.2 dup: SFARI genes OR=3.3, FDR=1.4×10^-13^; PCDH19: SFARI genes OR=3.1, FDR=1.5×10^-5^; Timothy syndrome: SFARI genes OR=2.0, FDR=4.3×10^-2^; 16p11.2 del: brain disorder genes OR=2.4, FDR=6.6×10^-3^; 16p11.2 dup: brain disorder genes OR=2.2, FDR=1.2×10^-2^; PCDH19: brain disorder genes OR=3.2, FDR=2.2×10^-3^; Fisher exact test, Figure 2g and Extended Data Figure 7a; Methods). Gene set enrichment analysis at this early time point found downregulation of terms related to neural precursor proliferation (e.g., “*Beta catenin binding*” “*Neural precursor cell proliferation*” and “*Canonical WNT signaling pathway*”) in all 4 of these forms (Extended Data Figure 7b and Supplementary table 4). These findings further demonstrate convergence during the early stages of neurogenesis across different genetic forms of ASD.

### Dysregulated biological processes and cell types

To identify biological pathways that were disrupted in ASD across development, we investigated whether there were any shared categories across the multiple mutational forms and across time. We observed that 6 out of the 9 different forms of ASD profiled were enriched for upregulated genes related to neuronal cell fate and protein translation-related terms (Figure 3a and Supplementary table 5) (Mohammad et al., 2021). We also observed broad down regulation of synapse related terms and ion channel related such as *“Synaptic membrane”* and *“Gated channel activity”*, across multiple timepoints and all the major mutational forms (Figure 3a and Supplementary table 5). To further explore the relationships of gene expression changes to specific cell types, we deconvolved bulk RNA sequencing into cell-type transcriptomes using single cell RNAseq from hCOs ((Khan et al., 2020); Extended Data Figure 8). This cell type specific analysis determined that both the least mature progenitors (CycProg) and more mature neuronal cell types (ExNeu-2) showed similar down-regulation across most conditions suggesting changes in the timing of neuronal maturation (Figure 3b and Extended Data Figure 9). A recent paper that examined 16p11.2 deletion in organoid models also found a decrease in cycling progenitors and an increase in neurons (Urresti et al., 2021).

**Figure 3.**
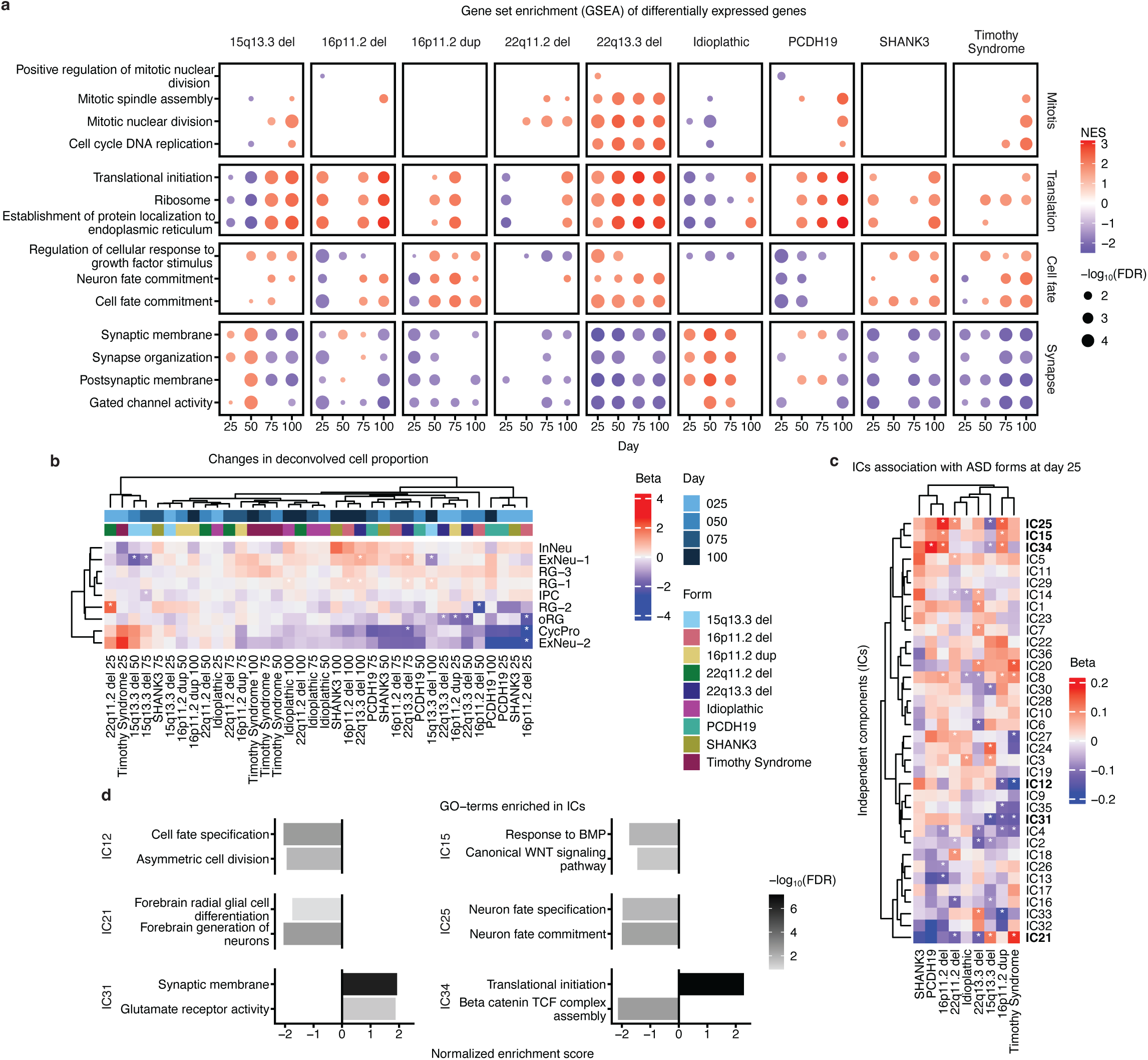
Pathways and cell types affected across multiple forms of ASD. (a) Select gene ontology (GO) terms enriched in upregulated (red) or downregulated (blue) genes in each of the ASD forms. Color corresponds to normalized enrichment score (NES). Point size corresponds to the level of significance (-log_10_(FDR)). GO-terms were categorized into four categories: Mitosis, Translation, Cell Fate and Synapse. (b) Hierarchal clustering of changes in cell proportions between the different forms of ASD and the controls. Top most annotation bar represents the differentiation day (shades of blue) and lower annotation bar represents the ASD form (colorful). CycPro - cycling progenitor, RG-1 - radial glia cluster 1 (early), RG-2 - radial glia cluster 2 (late), RG-3 – radial glial cluster 3, IPC - intermediate progenitor cells, ORG - outer radial glia, ExNeu-1 - excitatory neurons cluster 1 (early), ExNeu-2 - excitatory neurons cluster 2 (late), InNeu - Intermediate neurons. (c) Association of independent components (IC) with the different forms of ASD at day 25. (d) Select gene ontology (GO) terms enriched for the positively or negatively contributing genes for the indicated ICs. Negatively enriched terms are downregulated in ICs that are positively associated with forms of ASD and are upregulated in ICs that are negatively associated with forms of ASD. * FDR < 0.05, ** FDR < 0.01, *** FDR < 0.005.

Next, we performed independent components (IC) analysis to determine if these changes related to neurogenesis and neural differentiation were robustly identified using an orthogonal method. IC analysis is an unsupervised approach, which separates a multivariate signal into additive subcomponents, and allows for the uncovering of underlying biological information by exploring gene expression features (Methods). We observed that the ICs exhibited similar patterns to the expression-based clustering (Rand index= 0.58) with 16p11.2 deletion and PCDH19-related disorder and 16p11.2 duplication and Timothy syndrome clustering together (Figure 3c and Extended Data Figure 10). IC15, IC25, and IC34 were up regulated in both 16p11.2 deletion IC15 beta=0.10, FDR=0.01; IC25 beta=0.18, FDR=1.3×10^-3^; IC34 beta=0.14, FDR=2.3×10^-4^) and 16p11.2 duplication (IC15 beta=0.10, FDR=0.02; IC25 beta=0.14, FDR=0.02; IC34 beta=0.11, FDR=7.4×10x^-3^) and to a lesser degree with PCDH19 and Timothy syndrome. These up-regulated ICs were enriched for processes involved in neuronal precursor renewal and differentiation, such as “*Response to BMP*”, “*Beta catenin-TCF complex assembly*”, and “*Neuron fate specification*” (Ohtaka-Maruyama and Okado, 2015) (Figure 3d and Supplementary table 6) further pointing to change in the differentiation trajectory of early neurons. The role of these pathways in ASD has previously been suggested more widely (Jourdon et al., 2023; Kumar et al., 2019; Paulsen et al., 2022), highlighting their potential role as a point of convergence of several forms of ASD. Two other notable ICs, IC12 and IC31, which were up regulated for synaptic terms and down regulation for glial differentiation, were downregulated in 16p11.2 duplication (IC12 beta=-0.13, FDR=8.4×10^-3^; IC31 beta=-0.13, FDR=1.11×10^-3^) and Timothy syndrome (IC12 beta=-0.19, FDR=2.9 ×10^-3^; IC31 beta=-0.14, FDR=2.9 ×10^-3^_;_ Figure 3c-d and Supplementary table 6) similar to what was observed in the differential expression analysis. Taken together, transcriptional changes reflected by these terms suggests an asynchronous developmental trajectory, which is most pronounced at very early timepoints across multiple distinct genetic forms of ASD, most robustly detected in 16p11.2 deletion, 16p11.2 duplication, PCDH19 mutation and Timothy syndrome, consistent with other recent observations in 3 different forms of ASD (Paulsen et al., 2022).

In addition to the convergent changes seen across multiple forms, we saw some changes that were unique to specific forms. For example, 22q13.13 showed increases in “*mitotic nuclear division”* and “*cell cycle DNA replication”* across days. This increased mitotic activity is also found in the ICA analysis, as IC2, a component that is downregulated in mitosis terms, is also downregulated in 22q13.13 deletion (Figure 3c and Extended Data Figure 10; beta=–0.13, FDR=3.7 ×10^-2^). In the case of 15q13.3 deletion, we observed a shift in developmental patterns of gene expression, characterized by initial elevations in terms such as ‘Synapse Organization’ and ‘Synapse Membrane’ which were accompanied by simultaneous reductions in translation-related terms like ‘Ribosome’ and ‘Translation Initiation.’ Subsequently, there were later declines in synapse-related terms and an increase in translation-related terms, evident at both day 75 and day 100 (Figure 3a).

### Network analysis reveals convergence between different ASD forms

Since transcripts do not act independently, but as part of highly regulated transcriptional networks (Parikshak et al., 2015), we constructed gene networks to identify modules of highly co-expressed genes, a powerful way to identify biological processes in an unsupervised manner (Li et al., 2018; Oldham et al., 2008; Parikshak et al., 2015). We then determined whether they were associated with specific time points, mutational forms of ASD, or shared across distinct forms. Considering that the largest changes in expression occurred on day 25 (Figure 2d), we concentrated on modules derived from the samples at this time point (Figure 4, Extended data Figure 11 and Supplementary table 7). Day 25 of differentiation corresponds to a very early period in cortical development, consisting primarily of early cycling progenitors and radial glia (Kanton et al., 2019; Paşca et al., 2015). Notably, we observed that the 22 identified modules were based on co-expression relationships from day 25 data, they were all preserved at all other time points, suggesting that they represent robust, ongoing biological processes over this 100-day period. Moreover, their conservation in *in vivo* (Li et al., 2018) and *in vitro* (Adhya et al., 2021; Flaherty et al., 2019; Lin et al., 2016; Schafer et al., 2019) data sets (Extended Data Figure 12) further demonstrates their reproducibility and relationship to *in vivo* neurodevelopment, showing that these modules represent generalizable human neurodevelopmental processes.

**Figure 4.**
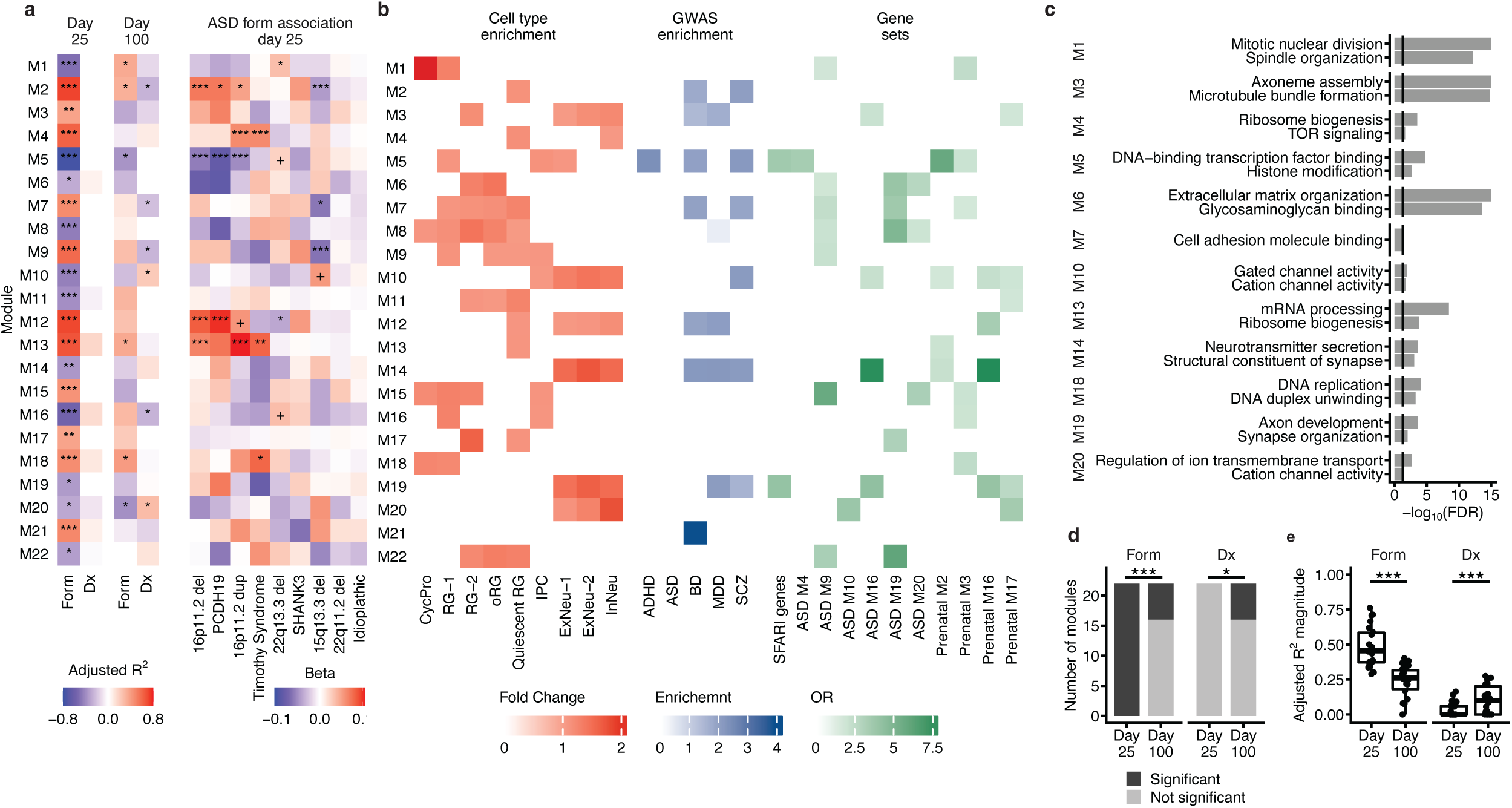
Network analysis of gene expression. (a) Adjusted R-squared of modules significantly associated with either a form of ASD (Form) or with the presence of ASD (Dx) at both day 25 and day 100 (left). Changes in the module eigengene (ME) in each of the forms of ASD compared to control (right). (b) Cell type enrichment of the modules using EWCE (red), GWAS enrichment using stratified LD score (blue) and select ASD related gene set enrichment using fisher test (green). (c) Select gene ontology (GO) terms enriched in modules. Black lines represent the FDR = 0.05 cutoff. (d) Number of modules associated with either ASD forms (specific mutations) or with ASD broadly (Dx). Significance was tested using Fisher’s exact test. (e) Magnitude of association of modules with either ASD forms or with ASD broadly (Dx). Significance was tested using Welch’s two sample t-test. * FDR < 0.05, ** FDR < 0.01, *** FDR < 0.005.

At the earliest timepoint, there were 22 modules that differentiated between any of the forms of ASD and controls. However, no single module was associated with all the different forms of ASD or differentiated ASD broadly from controls (Fig 4a). Eleven of the modules were significantly associated with at least one of the forms of ASD, and 5 were associated with multiple forms, indicative of convergence among those forms at day 25 (Figure 4a). These included M4, M5 and M13 which associated with two or more forms of ASD including 16p11.2 deletion, 16p11.2 duplication, Timothy Syndrome and PCDH19-related disorder (M4: 16p11.2 dup beta=0.06, FDR=3.3×10^-4^, Timothy Syndrome beta=0.06, FDR=3.8×10^-3^; M5: 16p11.2 del beta=-0.06, FDR=2.3×10^-6^, 16p11.2 dup beta=-0.05, FDR=5.6×10^-4^, PCDH19 beta=-0.08, FDR=1.7×10^-5^; M13: 16p11.2 del beta=0.07, FDR=4.0×10^-3^, 16p11.2 dup beta=0.11, FDR=7.4×10^-7^, Timothy Syndrome beta=0.08, FDR=6.6×10^-3^). Notably, we observed no modules associated with idiopathic ASD in our cohort, consistent with its heterogeneity. Pathway analysis revealed that these modules are involved in translation (M4, M13) and transcriptional regulation (M5) in radial glia, intermediate progenitors, and early neurons (Figure 4b-c and Supplementary table 8).

In contrast, at day 100, which corresponds to early mid-gestation (Gordon et al., 2021; Paşca et al., 2015; Trevino et al., 2020) and contains a large pool of post mitotic neurons and remaining radial glia, we identified only 6 modules that were associated with any of the various forms of ASD (Fisher test p=3.6×10^-7^; Methods; Figure 4d). This indicates that the differences between cases and controls become less pronounced with time, mirroring observations within differential expression. Further emphasizing this, the magnitude of association of the modules with the individual ASD forms was significantly lower at day 100 compared to day 25 (day 25=0.48, day 100=0.24, Welch two sample t-test: t_40.06_=6.3, p=1.7×10^-7^; Figure 4e). Conversely, at day 100 significantly more modules differentiated ASD broadly from controls (6 at day 100 compared to 0 at day 25; Fisher test p=0.02; Figure 4d) and the association of the modules with ASD broadly was significantly higher compared to day 25 (day 25=0.03, day 100=0.11, Welch two sample t_30.69_=–3.0, p=5.0×10^-3^; Figure 4e) suggesting that the different forms become more similar as differentiation progresses, analogous to what we previously observed using orthogonal analyses (Figure 2c).

To understand how these convergent modules might be related to core processes underlying ASD, we next asked whether any were enriched in genes harboring ASD associated genetic susceptibility variation (Methods). We found that two of these modules, M5 and M19, were significantly enriched for rare genetic risk variation associated with ASD (Figure 4b M5: OR=3.6, FDR= 9.4×10^-12^; M19: OR=4.1, FDR= 2.1×10^-6^). This indicates that known ASD risk genes are strongly co-expressed following shared perturbation of neural development in multiple rare forms of ASD, similar to what we and others observed in development *in vivo* (Liao et al., 2023; Parikshak et al., 2013; Walker et al., 2019; Willsey et al., 2013).

Most notable in this regard is M5, which was significantly downregulated across several different forms of ASD at day 25 specifically (adjusted R^2^=-0.76, FDR = 5.2×10^-15^; 16p11.2 deletion beta=-0.06, FDR=2.3 ×10^-6^; 16p11.2 duplication beta=-0.05, FDR=5.6 ×10^-4^; PCDH19 beta=- 0.08, FDR=1.7×10^-5^) (Figure 4a), as well as across the entire course of differentiation (16p11.2 deletion : t-value=-3.5, FDR=2.0×10^-3^; 16p11.2 duplication: t-value=-3.6, FDR=2.0×10^-3^; PCDH19: t-value=-2.6, FDR=2.0×10^-2^; SHANK3: t-value=-2.3, FDR=3.8×10^-2^; Methods; Extended Data Figure 13) suggesting that it plays a central role downstream of these 4 ASD-associated mutations throughout differentiation. M5’s relationship to *in vivo* human biology is supported by its significant overlap with two co-expression modules that peak in early to midfetal gestation human brain that were significantly enriched for rare ASD risk mutations ((Parikshak et al., 2013): Prenatal M2 OR = 5.8, FDR = 7.2×10^-32^; Prenatal M3 OR = 2.0, FDR = 2.9×10^-4^). Remarkably, we also observed that M5 was also enriched for genes co-expressed within a neuronal module that is downregulated in postmortem cerebral cortex from ASD subjects ((Parikshak et al., 2016); ASD M4, OR = 3.5, FDR = 3.3×10^-5^), providing further evidence that it represents convergent biology with relevance to processes occurring *in vivo* in ASD. Functionally, M5 was enriched for terms related to regulation of gene expression such as “*Histone modification*” and “*DNA-binding transcription factor binding*” (Figure 4c and Supplementary table 8) and was enriched in developing cell types: early radial glia (RG-1; fold change=1.1, FDR = 4.5×10^-6^), intermediate progenitors (IPCs; fold change=1.1, FDR = 4.5×10^-6^), and early cortical neurons (ExNeu-1; fold change=1.1, FDR = 3.0×10^-5^; Figure 4b).

M19, the second module which was enriched for rare genetic variation associated with ASD, was also down-regulated across several ASD forms (day 25 adjusted R^2^=-0.35, FDR = 0.01) (Figure 4a-b). Similar to M5, M19 also showed enrichment for rare ASD risk genes (SFARI genes, Figure 4b). M19 also overlapped with two later expressed, neuronal, fetal cortical co-expression modules that were significantly enriched for ASD risk ((Parikshak et al., 2013):Prenatal M16 OR = 4.1, FDR = 1.7×10^-5^; Prenatal M17 OR = 2.7, FDR = 1.2×10^-3^). Functionally, M19 is annotated as a synaptic neuronal module, evidenced by its enrichment for terms related to synaptic function such as “*Synapse organization*” and “*Neurotransmitter secretion*”, as well as being enriched for all neuronal cell types (ExNeu-1 fold change=1.5, FDR=3.0×10^-6^; ExNeu-2 fold change=1.6, FDR=3.0×10^-6^; InNeu fold change=1.5, FDR=3.0×10^-6^).

### M5 is an upstream regulator of ASD risk genes

Since the early expressed ASD risk-gene containing module, M5, was significantly enriched for terms related to regulation of gene expression (Figure 4c), we tested the regulatory relationships between M5 and other modules associated with various forms of ASD. We examined the enrichment of motifs upstream of each of the modules and annotated them to high confidence transcription factor binding sites (Figure 5a and Supplementary table 9; Methods). We then determined the hierarchical regulatory relationships among modules (Methods; (Verfaillie et al., 2015)), finding that M5 and M1 contained the highest levels of predicted up-stream transcriptional regulators of other modules (Figure 5c). Transcriptional targets of M5 were enriched in 10 of the ASD-associated modules, among which was M19, another module enriched for ASD risk genes (Figure 5b and Figure 4b). This highlighted M5’s potential role as a causal driver of the changes seen in ASD, which is consistent with its expression at early stages (Extended Data Figure 13, and (Parikshak et al., 2013)).

**Figure 5.**
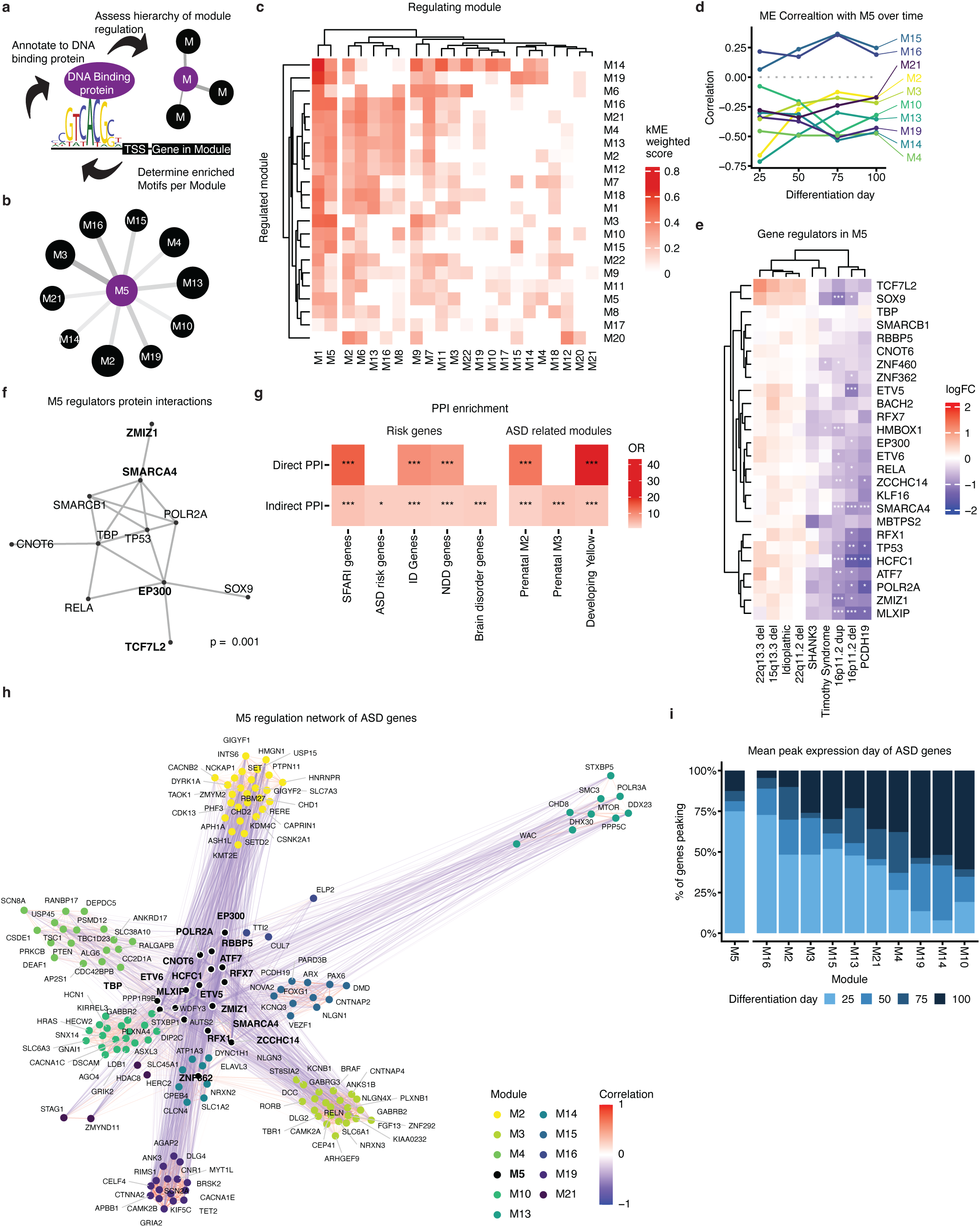
Module regulatory relationships reveal a key upstream regulator module. (a) Schema showing the identification of enriched motifs workflow using RcisTarget. Enriched motifs upstream of module gene lists are annotated to high confidence DNA binding proteins. (b) Modules regulated by M5. Line width represents the kME weighted score and circle size represents proportion of genes regulated by M5 genes. (c) Correlation of module 5 eigengenes with downstream modules across differentiation, showing negative correlations with most downstream modules across time. (d) Heatmap of log fold-change (logFC) of gene regulators in M5 at day 25 shows strong decreases across several forms of ASD. (e) Protein-protein interactions of the regulators in M5 genes. These genes form a strongly significant protein-protein interaction (PPI) network. (f) Enrichment of the regulators in M5 as well as their indirect network for ASD genes from either SFARI or from large scale whole exome (WES) sequencing studies (Ruzzo et al., 2019a; Satterstrom et al., 2020) as well as with neurodevelopmental disorders (NDDs) and intellectual disability (ID) risk genes (Leblond et al., 2021) (g) Correlation of the transcriptional regulators in M5 with ASD risk genes found in module downstream of M5. M5 genes are in bold and marked with a black ring. The overall majority of the correlations were negative (blue) whereas the genes within modules were positively correlated (red). Top GO-terms for the ASD genes in each module are presented. (h) Peak expression day of the regulators in M5 and the ASD genes that they regulate. Color bars represent the percent of genes whose mean expression peak in the corresponding differentiation day. * FDR < 0.05, ** FDR < 0.01, *** FDR < 0.005.

Other modules downstream of M5 were associated with various processes known to be dysregulated in neurodevelopmental disorders (NDDs) (Figure 5b,c and Figure 4c). We observed a relationship between M5 down-regulation and the up-regulated expression of genes in its downstream putative target modules, suggesting that M5 may be negatively regulating these modules. This was further supported by the negative correlation seen between the M5 ME and the majority (8/10) of the downstream modules MEs across differentiation (Figure 5d). Since genes in a module can vary in their degree of module membership (Langfelder and Horvath, 2008), we next asked if the transcriptional regulators (i.e., the transcription factors and chromatin modifiers) within M5 followed the same down-regulation pattern seen across all genes in the M5 module overall. Indeed 65% (17/26) of the annotated transcriptional regulators (Methods) in M5 were significantly down-regulated in at least one of the forms of ASD with most (8/9) of the other regulators trending in the same direction (Figure 5e).

To further elaborate the relationship between M5 regulatory targets and ASD, we asked whether genes regulated by M5 drivers in addition to those in M19, were also enriched for ASD genetic risk factors. Indeed, even though the individual co-expression modules containing these genes were not enriched for ASD risk genes, we found that the M5 regulated genes within these other M5 regulated modules were enriched for ASD risk genes (OR=2.0, p =3.6×10^-9^). A total of 261 SFARI 1,2, or S genes within enriched downstream modules were predicted to be regulated by at least one M5 regulator (Figure 5h). While almost all of the M5 regulators are predicted to modulate expression of multiple SFARI genes (24/26), a few have targets that are notably enriched in ASD risk genes including *ZNF460* (22% of predicted targets in enriched modules are SFARI genes), which is predicted to regulate genes including *NLGN2, POGZ, SATB1,* and *SETD5* and M5 regulator *ZCCHC14* (17% of predicted targets in enriched modules are SFARI genes) which regulates *ASH1L1, DYRK1A, and KMT2E* among others. Many of the SFARI genes are regulated by multiple M5 regulators; among the most convergently targeted are *DYRK1A* (target of 8 M5 regulators), *POGZ* (target of 7 M5 regulators), *SATB1* (target of 7 M5 regulators), and *SETD5* (target of 7 M5 regulators). Noticeably, these ASD risk genes were mostly (68%) negatively correlated with the M5 regulators (Figure 5h). Consistent with their role as transcriptional regulators, the driver genes within M5 peak in expression earlier than the ASD genes they are regulating (Figure 5i and Extended Data Figure 13). These M5 regulated ASD risk genes included many high confidence risk genes associated with synapse-related function, such as *CNTNAP2* and *NLGN1* (M16), forebrain neuron differentiation, such as *FOXG1* and *PAX6* (M15), and epigenetic remodelers including *SET*, *DYRK1A*, *KMT2E*, and *CHD2* (M2) (Figure 5h), which are all processes previously related to ASD (Gilbert and Man, 2017; Parikshak et al., 2013; Satterstrom et al., 2020). Interestingly, these genes also included *PCDH19* (M15) and *CACNA1C*, the gene mutated in Timothy syndrome (M10) (Figure 5h), further reflecting this transcriptional network’s role as point of convergence in ASD. Taken together, these findings suggest that a complex of transcriptional regulators orchestrates distinct biological pathways related to neurogenesis and neuronal maturation altered across ASD through regulatory relationships occurring during early stages of brain development.

### M5 regulators form a protein-protein interaction (PPI) network enriched for ASD risk genes

To determine if these transcriptional regulators represent some generalizable, known biological processes, we asked if they were more likely to interact at the protein level, finding that indeed they are predicted to form a highly significant protein-protein interaction (PPI) network (p = 0.001; Figure 5f). The PPI network represented by module M5 genes was enriched for transcription factor regulation and chromatin binding related GO terms (Supplementary table 10). Further examination of this PPI network also indicated that it was significantly enriched in known ASD risk genes, as well as in risk for other neurodevelopmental disorders (direct PPI - SFARI 1,2,S genes: OR=15, FDR=2.0×10^-4^; NDD; OR=8.4, FDR=3.7×10^-4^; ID; OR=9.7, FDR=2.0×10^-4^; Indirect PPI - SFARI genes: OR=3.7, FDR=4.6×10^-11^; ASD risk genes: OR=2.4, FDR=3.0×10^-2^; NDD genes: OR=2.1, FDR=1.7×10^-7^; ID Genes: OR=2.4, FDR=1.9×10^-9^; brain disorder genes: OR=2.4, FDR=3.6×10^-3^; Fig 5g, Methods). Moreover, similar to the transcriptome-based M5, the PPI network strongly overlapped with the early to mid-gestation brain co-expression networks that have previously been associated with ASD ((Parikshak et al., 2013): Direct PPI - Prenatal M2: OR=12.3, FDR=2.5×10^-4^), including common variant-enriched modules (Walker et al., 2019): Direct PPI - Developing Yellow: OR=43, FDR=7.1×10^-10^).

We next tested whether the predicted PPI interactions among M5 genes based on public databases (Figure 5f), were observed in human neural progenitor cells (hNPCs), given that protein interactions can be highly cell-type specific and most available data is not from neural cell-types (Pintacuda, 2023). We tested antibodies against putative TFs or chromatin regulatory proteins within the predicted PPI and conducted immunoprecipitation followed by mass spectrometry (IP-MS) on proteins with reliable signal (Figure 6a-b; Extended Data Figure 14a; Methods). We then examined whether we could validate the predicted PPI by examining connections between M5 proteins (Methods). We found that M5 regulators with reliable IP-MS signal do indeed form a significant PPI with each other and with a subset of the predicted gene lists (Figure 6c-f), including SFARI genes (score 1,2, and S combined and score S; Figure6c,e), NDD genes ((Leblond et al., 2021); Figure 6c,e, Extended Data Figure 14b-d) and genes enriched in DD as well as NDD genes not enriched in DD or ASD (Fu et al., 2022), validating database predictions (Figure 5e). Of the 5 M5 regulators that passed QC (methods), we found that most were connected to 3/4 of their possible connections, except for *POLR2A* (Figure 6d). This PPI includes BAF complex members *SMARCB1*(BAF47) and *SMARCA4* (BRG1) as well as *EP300*, an important coactivator known to interact with BAF complex members (Blümli et al., 2021), which play critical roles during neurogenesis and mammalian cortical development (Braun et al., 2021; Jin et al., 2022; Ninkovic et al., 2013; Son and Crabtree, 2014; Tuoc et al., 2013) and *TBP*. These findings show that M5 regulators interact with a network of genes essential for normal neurodevelopment, both in terms of predicted transcriptional control and direct protein-protein interactions in neural progenitors.

**Figure 6.**
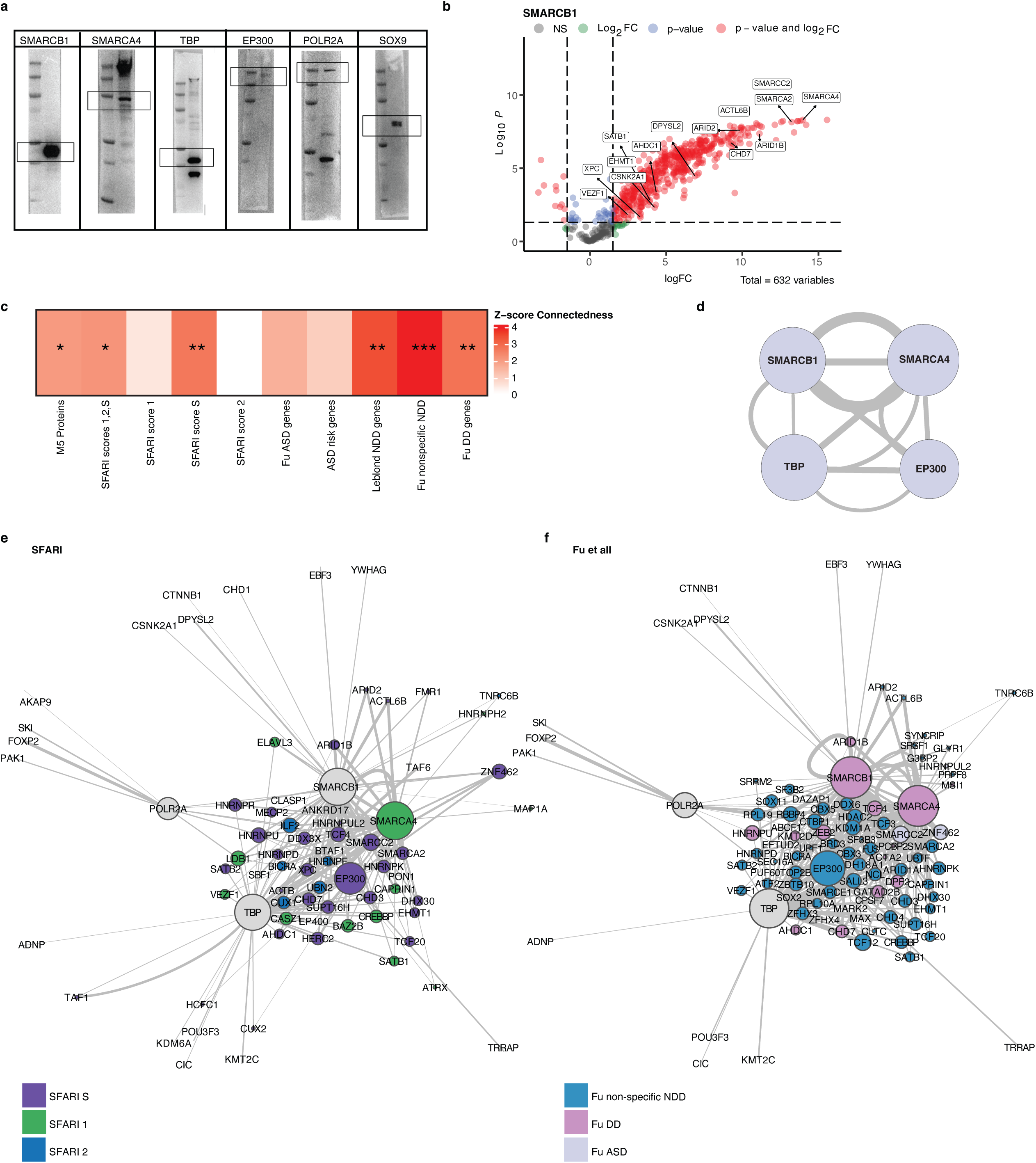
M5 regulators form a protein-protein interaction (PPI) network enriched for neurodevelopmental risk. (a) Western blots confirm antibody binding for proteins of interest. (b) Example volcano plot shows significantly bound proteins. Highlighted proteins are SFARI risk genes. (c) Z-score connectedness of gene-lists show that M5 proteins detected in IP-MS experiments form a significant PPI and that this network is significantly connected with other neurodevelopmental risk such as SFARI genes (1,2,S), NDD genes and Autism genes (Leblond et al., 2021; Ruzzo et al., 2019b; Satterstrom et al., 2020) and genes enriched for DD, ASD, or NDD broadly (Fu et al., 2022). d-f) Connectivity networks of key neurodevelopmental gene lists. Size of circle represents number of connections, edge width represents logFC between nodes. Color represents gene-list. Grey nodes are network baits not in featured gene-list. * FDR < 0.05, ** FDR < 0.01, *** FDR < 0.005

### M5 regulators regulate their predicted targets

Having validated the PPI network in developing neurons, we next sought to experimentally validate the regulatory role of M5 transcription factors and chromatin regulators on their predicted downstream targets. We leveraged the ability to perform a pooled, CRISPR/Cas9 knockdown screen to assess downstream gene expression on a single cell basis (Methods; (Datlinger et al., 2017))(Figure 7a, Extended Data Figure 15). We applied this to neural cells derived by NGN2 induction in hiPS cells (Methods) that rapidly generates large numbers of relatively homogeneous neural progenitor cells (hNPC). We targeted the promotors of 26 M5 putative transcriptional regulators in hNPCs using 3 guides per gene ((Guss et al., 2024); Methods) and were able to achieve consistent downregulation (Figure 7b, Extended Data Figure 15), ranging from ∼25% (*POLR2A*) to ∼90% decrease (*ZNF460*) with 80% of the genes demonstrating a reduction in expression of 50% or more (Figure 7b). We observed that hNPCs separated into two classes, based on those that were mitotically active (NPCs showing consistent G2 mitosis markers such as *TOP2A*, *CDK1*, (termed cNPCs), Extended Data Figure15) or those not expressing these markers (termed hNPCs). For most targets, knockdown did not have a major impact on survival or proliferation, with 24/26 knockdown targets showing no difference in proportion of cells compared to controls except for *SMARCB1* and *TP53*, which both showed increased proportions of both G2/mitotic and hNPCs (Extended Data Figure 15).

**Figure 7.**
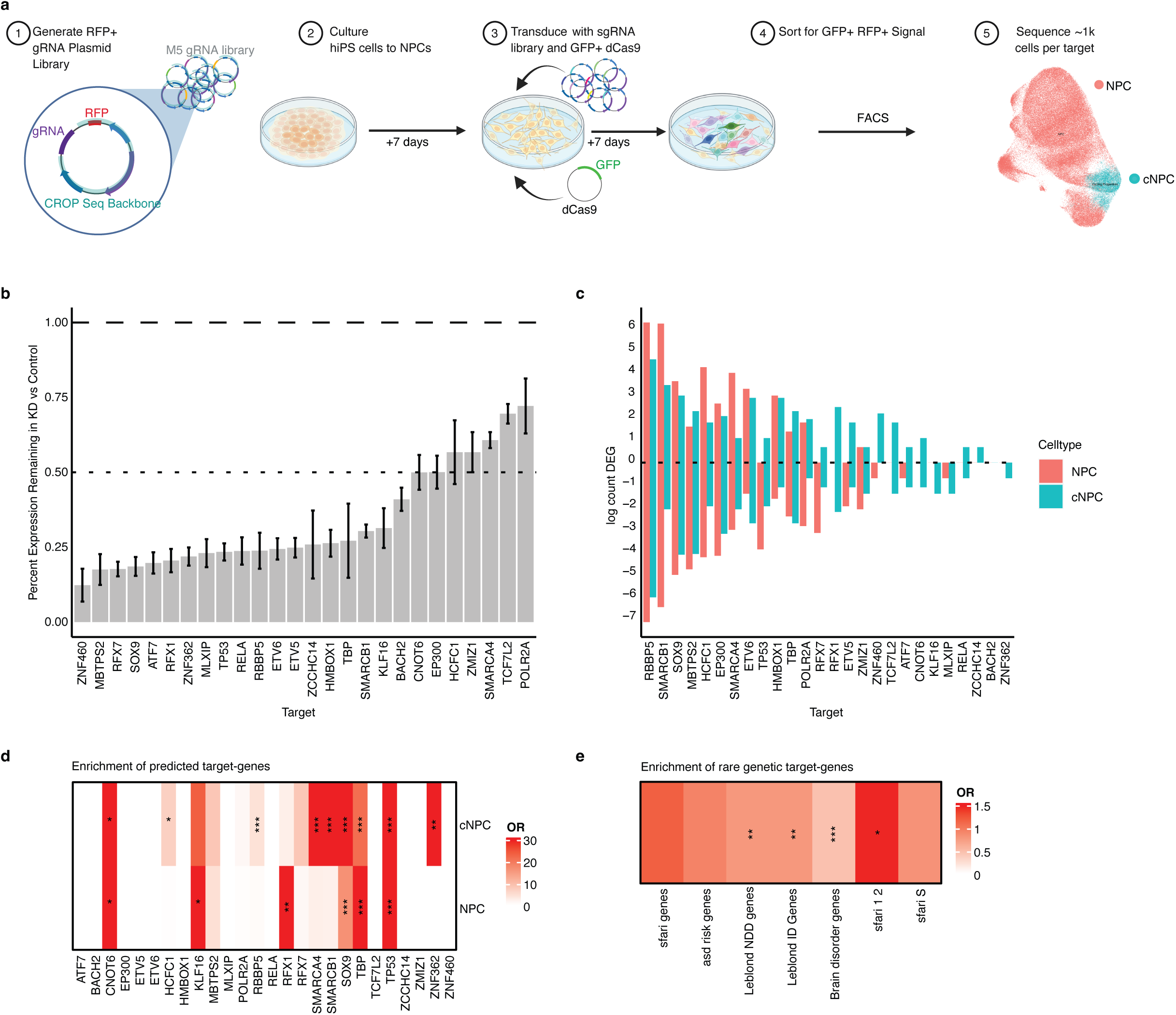
CRISPRi of M5 regulator genes shows effect on predicted targets and genes linked to Neurodevelopmental disorders. (a) Schematic of experimental approach. 117 gRNAs corresponding to 26 genes and 39 NTC were designed and transduced into SNaP’s. Seven days post transduction, single cells were sorted by Fluorescent Activated Cell Sorting (FACs) and single cell RNA seq was performed (n=6 replicates). NPCs were separated into cycling neural progenitors (cNPCs) and Neural progenitor cells (NPCs). (b) Mean ratio of target gene expression (y-axis) remaining in targeted cells (x-axis) compared to NTC. (c) Log counts of differentially expressed genes (DEGs) (p value adjusted <.05) in each target. (d) Enrichment of predicted regulated genes in DEG dataset. (e) Enrichment of DEGs in rare genetic variation sfari genes (1,2,S), asd risk genes, neurodevelopmental disorder genes, intellectual disability risk genes, and brain disorder genes (Leblond et al., 2021; Ruzzo et al., 2019b; Satterstrom et al., 2020). * FDR < 0.05, ** FDR < 0.01, *** FDR < 0.005

*TP53* knockdown-induced increased proliferation is consistent with its well-known role as a tumor suppressor (Phan et al., 2022). *SMARCB1* loss is also associated with tumorigenesis in patients, though previous studies have not seen an increased proliferation phenotype in NPCs (Parisian et al., 2020). Cells expressing gRNAs targeting *POLR2A* showed only ∼25% knockdown, so although *POLR2A* gRNA-expressing cells did not show a significant difference from controls, this could be due to a limit on the amount of *POLR2A* knockdown tolerated by cells (Yu et al., 2019).

We observed a wide spectrum of magnitude of differential expression following pooled CRISPRi, with *RBBP5* knockdown leading to >1250 downregulated and >500 upregulated genes in NPCs to genes such as *RELA* and *BACH2* where we observed no significant differential expression (Figure 7c). *RBBP5* is part of the WRAD complex, which supports H3K4 methylation for SET-domain containing methyltransferases, many of which are major neurodevelopmental risk genes (Ernst and Vakoc, 2012). *RBBP5* knockdown led to massive changes in gene expression including SWI/SNFI member *SMARCA1* and synapse genes *DLGAP1, NRXN3,* and *CACNA1C*, the gene mutated in Timothy syndrome. Not every M5 regulator showed strong evidence for recapitulating phenotypes seen in hCOs, but knockdown of most M5 members produced a transcriptional response that was correlated with changes seen in organoids (Extended Data Figure 16). Overall, knockdown of 11 of the 26 M5 regulatory drivers showed significant differential expression enrichment of their predicted targets - 9/26 in cNPC and 6/26 in hNPCs, confirming their regulatory relationship (Methods, Figure 7d).

We also examined whether genes that are associated with rare genetic variation in NDDs were enriched in differentially expressed genes downstream of the M5 regulators. Parallel with the earlier bio-informatic predictions, we found that genes associated with NDDs and intellectual disability in addition to ASD risk genes (Leblond et al., 2021; Ruzzo et al., 2019b; Satterstrom et al., 2020)(SFARI levels 1 and 2) were enriched in DEGs (Figure 7e). Pathways downstream of several of the individual M5 regulators, including *SMARCB1*, *SOX9*, and *RBBP5* lead to disruptions to core neurodevelopmental pathways (Extended Data Figure 17), recapitulating their effect in organoid models (Figure 3a). For example, knockdown of *EP300* led to decreases in axon generation related terms, while knockdown of *SMARCB1* led to broad changes including decreases in proliferation, axon generation, and synapse maturation-related terms (Extended Data Figure 17). We find decreases in *SOX9* led to both decreased neural fate commitment terms such as “Neural precursor cell proliferation” and synaptic terms encompassing early-expressed synaptic genes such as “Synapse assembly”. Synapse-related genes affected by *SOX9* knockdown include *NRXN3, GRIN2B, and NLGN3,* demonstrating impact on synapse-expressed genes even prior to the presence of synapses. While examining individual cellular responses to each M5 regulator is important to validate predictions of regulation, future studies should examine the timing of decreases of the regulators in organoid models to better understand how changes in these genes might alter cellular transitions during brain development.

## Discussion

Here we characterize the biological impact of many different genetic forms of ASD modeled in a large, unprecedented cohort of hCO, based on quantitative analyses rooted in genome-wide transcriptomic profiling. We identified robust clusters of different ASD forms spanning across stages of cerebral cortical development, suggesting several distinct developmental trajectories. Orthogonal analytic approaches showed that as time progressed these trajectories converged, consistent with canalization of development where changes in the early stages of brain development are buffered as development progresses (Wagner et al., 1997). In general, these trajectories converge on decreased neuronal maturation across timepoints, with most forms showing changing trajectories starting from day 25 and all, except for idiopathic ASD. While the alterations in maturation are convergent, the path to these changes seems to be distinct depending on different mutations. This is consistent with previous observations in a cohort profiling three other genetic forms of ASD (Paulsen et al., 2022). Several individual genetic forms showed opposing effects. For example, PCDH19-related disorder and 16p11.2 deletions showed diminished neural fate commitment, which single cell deconvolution confirmed reflects a diminished progenitor pool. In contrast, 22q13.3 deletion showed patterns consistent with an over-proliferation of the neural progenitor pool. Therefore, altered developmental trajectories seen here are not as simple as ‘delayed’ or ‘accelerated’ but rather point to specific cellular transition states, such as early maturing neurons, as times of convergent vulnerability. We note that these are still relatively early stages of corticogenesis in hCO, before gliogenesis and synaptogenesis have started; at later stages these phenotypes may either be amplified or compensated for. This does not preclude early role of synaptic genes at pre-synaptic stages as it has been recently described for a number of disease relevant genes (Panagiotakos and Pasca, 2022).

We also describe a core transcriptional network of driver genes acting at early stages and identified their downstream target biological processes. This core set of convergent genes is changed across several distinct genetic forms and is enriched in rare and common genetic risk for ASD. Interestingly, M5 dysregulation may not be limited to ASD, as it was also enriched for common variation associated with multiple psychiatric disorders, reflecting the role of early neuronal differentiation in broad risk for psychiatric disorders (2019; Schork et al., 2019). These gene regulatory relationships, which we show form interacting protein complexes, connect ASD risk genes across different pathways, including WNT signaling, translation and mTOR pathways, and synapse formation into a convergent network. It is also worth noting that we identified few consistent changes in lines from idiopathic ASD. This is likely due to the substantial polygenicity and heterogeneity in ASD cases not harboring major risk mutations (Eyring and Geschwind, 2021). It is also interesting to consider that the effects of common variation may also impact later developmental stages than the early impact of transcriptional regulators and chromatin modifiers impacted by rare protein disrupting variants (Satterstrom et al., 2020) (Parikshak et al., 2013) (Leblond et al., 2021) (Fu et al., 2022).

This work and other recent work (Paulsen et al., 2022) (Li et al., 2023) demonstrates the utility of studying multiple mutations in parallel across developmental time to identified shared and distinct pathways in ASD. By describing a core transcriptional network of driver genes, known to be causal and identifying their downstream targets, which are also enriched in ASD risk genes, these data connect ASD genetic risk to convergent biological processes that occur during early neurodevelopment.

## Methods

### Collection and characterization of hiPS cell lines

Control hiPS cell lines in this study were reported and have been extensively characterized using standardized methods (Khan et al., 2020; Paşca et al., 2015; Yoon et al., 2019). hiPS cell lines from individuals with 11 idiopathic ASD s were obtained from Coriell. 22q11.2 deletion ((Khan et al., 2020), 22q13.3 deletion (Miura et al., 2020) and Timothy syndrome (Birey et al., 2022) hiPS cell lines were reported and characterized before. To confirm the various genetic mutations, whole genome sequencing was performed. Cultures were regularly tested for Mycoplasma and maintained mycoplasma free. Approval for reprogramming and differentiation of these cell lines was obtained from the Stanford IRB panel, and informed consent was obtained from all individuals.

### Reprogramming and hiPS cell culture

Human fibroblast cells were reprogrammed using the Sendai virus-based CytoTune iPS 2.0 kit (CytoTune 2.0, Invitrogen, A16517). Fibroblasts were plated in six-well plates two days before transduction to achieve a density of 2 × 10^5^ to 3 × 10^5^ cells per well. Cells were then transduced with the indicated set of Sendai vectors at a multiplicity of infection of 5–5–3 (*KOS*–*Myc*–*Klf4*) as we previously described (Khan et al., 2020). Virus was removed after 24 h and cells were cultured in fibroblast medium with 10% FBS (Gibco, 16000-044) in Dulbecco’s modified Eagle medium (DMEM, Gibco, 10569-010) for a total of seven days, with media changes every other day. On day 7 after transduction, cells were collected with TrypLE (Gibco, 16563-029) and plated in the fibroblast medium onto truncated recombinant human vitronectin (VTN-N, Gibco, A14700)-coated plates. The following day, fibroblast medium was changed to Essential 8 medium (Gibco, A1517001) and cells were fed daily until days 20–22. For further culture, individual colonies were manually dissected and transferred to fresh vitronectin-coated dishes. Once hiPS cell clones were established, they were routinely cultured on vitronectin-coated dishes in Essential 8 medium, and cells were split every 4–5 days using 0.5 mM EDTA (Invitrogen, 15575-020). hiPS cell clones/lines were also generated using retroviruses or non-integrating episomal vectors from fibroblasts (Bhaduri et al., 2020). We did not observe any systematic differences between cells reprogrammed by the two methods. Cultures were regularly tested for mycoplasma and maintained mycoplasma free.

hiPS cells were cultured as previously described (Yoon et al., 2019). Briefly, hiPS cells were cultured on six-well plates coated with recombinant human vitronectin (VTN–N, Gibco, A14700) in Essential 8 medium (Gibco, A1517001) and passaged with 0.5 mM EDTA (Gibco, 15575). To coat the six-well plates, 1 ml of vitronectin (diluted at a 1:100 ratio with Dulbecco’s phosphate-buffered saline (DPBS); Gibco, 14190) was added per well and then incubated at room temperature for 1 h. To passage hiPS cells with 80–90% confluency, cells were rinsed with 3–4 ml of DPBS per well, and then 1 ml of 0.5 mM EDTA (Gibco, 15575) was added and incubated for 7 min at room temperature. After the EDTA was removed, 2 ml of pre-warmed complete Essential 8 medium was added to collect cells. The cell suspension was then diluted in Essential 8 medium (1:6–1:20 depending on the hiPS cell line) and distributed on vitronectin-coated wells.

### Generation of hCOs from hiPS cells

For the generation of 3D spheroids, hiPS cells were incubated with Accutase (Innovate Cell Technologies, AT-104) at 37°C for 7 min and dissociated into single cells. To obtain uniformly sized spheroids, we used AggreWell 800 plates (STEMCELL Technologies, 34811) containing 300 microwells. Aproximately 3 x 10^6^ single cells per well in Essential 8 medium supplemented with the ROCK inhibitor Y-27632 (10 μM, Selleckchem, S1049), centrifuged at 100 g for 3 min to capture the cells in the microwells and incubated at 37°C with 5% CO_2_. After 24 h, day 0 of differentiation, spheroids were collected from each microwell by firmly pipetting (with a cut end of a P1000 tip) medium in the well up and down and transferring it into ultra-low-attachment plastic dishes (Corning, 3262) in Essential 6 medium (Gibco, A1516401) supplemented with two SMAD pathway inhibitors – dorsomorphin (2.5 μM, Sigma-Aldrich, P5499) and SB-431542 (10 μm, Tocris, 1614) together with Wnt pathway inhibitor XAV-939 (1.25 μM, Tocris, 3748). Media changes are performed daily, except for day 1. On the sixth day in suspension, neural spheroids were transferred to neural medium containing Neurobasal-A (Gibco, 10888), B-27 supplement without vitamin A (Gibco, 12587), GlutaMax (1:100, Gibco, 35050) and penicillin and streptomycin (1:100, Gibco, 15140). The neural medium was supplemented with epidermal growth factor (EGF, 20 ng ml^-1^, R&D Systems, 236-EG) and basic fibroblast growth factor (bFGF, 20 ng ml^-1^, R&D Systems, 233-FB) until day 24. From day 25 to day 42, to promote differentiation of the neural progenitors into neurons, the neural medium was supplemented with brain-derived neurotrophic factor (BDNF, 20 ng ml^-1^, Peprotech, 450-02) and neurotrophin 3 (NT3, 20 ng ml^-1^, Peprotech, 450-03), with medium changes every other day. From day 43 onward, only neural medium without growth factors was used for medium changes every 4-5 days.

### RNA-seq processing

Sequencing libraries were prepared using Truseq stranded RNA RiboZero Gold (Illumina) on ribosomal RNA depleted (RiboZero Gold, Illumina) RNA. The libraries were sequenced with 100-base-pair paired end reads on an Illumina HiSeq 4000. The reads were mapped to the human genome (hg38) with Gencode v25 annotations using STAR (v.2.5.2b) (Dobin et al., 2013) and gene expression was quantified using rsem (v1.3.0) (Li and Dewey, 2011). Genes that were expressed at very low levels (less that 10 reads in 30% of samples from a given day) were removed from the analysis. Samples with standardized sample network connectivity Z scores below –2 in each mutation were defined as outliers and removed (Oldham et al., 2012). To control for technical variation due to the sequencing and library prep we calculated the principal components of the Picard sequencing metrics (http://broadinstitute.github.io/picard/) using the CollectAlignmentSummaryMetrics, CollectRnaSeqMetrics, and MarkDuplicates modules and included them in our model. To infer genetic ancestry, we called SNPs from the aligned reads using the GATK (v3.3) Haplotype caller (McKenna et al., 2010). Sites with more than 5% missing samples, with rare minor allele frequency (less than 0.05) and Hardy– Weinberg disequilibrium (less than 1 × 10–6) were removed using plink (v1.09) (Purcell et al., 2007). We then used the remaining high-quality SNPs were used to run MDS together with HapMap3.3 (hg38) (International HapMap et al., 2010). The first two MDS values, referred to as genetic ancestry PC1/2, were then included in our model.

Sample identity was verified using the identity by descent (IBD) algorithm from PLINK (1.09) (Purcell et al., 2007). IBD was calculated for each pair of samples based on genotypes derived from RNA sequencing analysis as well as for all pairs of RNA and DNA sequencing samples. Samples with a 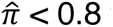 from other samples derived from the same individual were removed.

Sex was verified for each sample by detection of genes expressed by the Y chromosome. Sample with high levels of duplication (Picard tools: PERCENT_DUPLICATION > 65), high levels of intergenic mapping (Picard tools: PCT_INTERGENIC_BASES > 60) or with low levels of mRNA (Picard tools: PCT_MRNA_BASES < 0.5) were removed. All RNA-seq samples passing quality control are listed in Supplementary Table 1. Variance partitioning was performed using the variancePartition package (v1.20.0) with default parameters. Gene expression reproducibility was measured using Spearman’s correlation between every pair of samples in a given timepoint and mutation both within and between individuals.

### Whole Genome sequencing

Libraries were prepared using TruSeq DNA PCR-Free (Illumina) and were sequenced with 150-base-pair paired end reads on an Illumina HiSeq 4000. Sequencing data was processed as previously described (Ruzzo et al., 2019a).

The sequencing reads were mapped to the human genome (hg38) using Burrows-Wheeler Aligner (bwa-mem, v0.7.17) (Li and Durbin, 2009). The resulting bam files were merged using bamtools (v2.5.1) (Barnett et al., 2011) resulting in a single BAM file per sample. Duplicate reads were marked using the Picard MarkDuplicates tool (v2.5.0). Reads around indels were locally realignment using GATK’s IndelRealigner (v3.3). Base quality score was recalibrated using GATK (McKenna et al., 2010). Variant and non-variant bases in the genome were identified using GATK’s HaplotypeCaller (v3.3). Resulting VCFs were combined using GATK’s combineGVCFs (v3.3) and variants were called per chromosome using GATK’s GenotypeGVCFs (v3.3). Well-calibrated quality scores were generated using GATK’s Variant Quality Score Recalibration (VQSR; v3.3). Large chromosomal changes, resulting from iPSC chromosomal instability, were identified using DELLY (v0.8.7) (Rausch et al., 2012). Lines with large indels (>750Kb) that were confirmed using allele frequency in the RNA sequencing data and were removed from the analysis. This included 5 individuals where all lines were dropped (4 because of incorrect or absent mutations, 1 that did not pass QC). Twenty six lines in total (19 individuals) were dropped due to QC failures. These include 7 lines that were dropped due to failure to confirm mutation (4 individuals), 9 lines dropped due to large CNVs (7 individuals), 2 lines dropped due to leftover sendai virus (2 individuals), 4 due to additional or improper deletions (4 individuals), and 4 that were not fully characterized or did not pass QC (2 individuals). Point mutations in SHANK3, PCDH19 and CACNA1C were visually confirmed using Integrative Genomics Viewer (IGV; v2.9.4) (Robinson et al., 2011). Sequencing depth was calculated using Samtools depth command (v1.9) (Danecek et al., 2021). All WGS samples are listed in Supplementary Table 2.

### Differential expression

Gene counts were normalized using trimmed mean of M-values (TMM) (Robinson and Oshlack, 2010) as implemented in the calcNormFactors function from the edgeR package (v3.26.8) (Robinson et al., 2010). The mixed linear model used included differentiation day, sex, batch, the first two PCs calculated for genetic ancestry and 15 sequencings PCs (SeqPCs) as fixed effect and accounted for multiple samples coming from the same individual, hiPS cell line and differentiation and for samples that were sequenced multiple times as random effects. The model was implemented using the dream function from the variancePartition package (v1.20.0) (Hoffman and Roussos, 2021). Genes with FDR < 0.05 were considered to be differentially expressed.

### Independent component analysis

Independent component analysis (ICA) was performed using the JADE algorithm as implemented in the MineICA package (v1.30.0) (Biton et al., 2014). The number of components was set to 36 with 10,000 maximum iterations. Association of the IC with the forms of ASD was tested using a mixed linear model. The model was implemented using the lme function from the nlme package (v3.1.152). Contrasts were used to compare each form of ASD at each time point to the control at the same time point using the glht function from the multcomp package (v1.4.18). Like other component analyses IC directions are random and so enrichment was performed for both positively and negatively contributing genes. Unlike PCs, IC are not inherently ordered and were therefore ordered by the amount of kurtosis in each IC.

### Gene set enrichment analysis

GSEA was performed using the fgsea package (v1.10.1) (Korotkevich et al., 2019) on all genes with 1,000,000 permutations and a set size ranging from 30 to 500. Gene ontology gene sets (v7.0) were downloaded from http://software.broadinstitute.org/gsea/msigdb/. For differential expression genes were ranked by log2 fold-change and for ICA genes were ranked by the projection values of the genes on each component. Enrichment for high confidence ASD genes (gene score < 2 or syndromic genes) from the SFARI database (https://gene.sfari.org/database/gene-scoring/), from large scale ASD sequencing studies (Ruzzo et al., 2019a; Satterstrom et al., 2020), and risk genes for neurodevelopmental disorders (NDDs), intellectual disability (ID) and brain disorders (Leblond et al., 2021) was performed using Fisher’s exact test.

### Network analysis

Gene counts were normalized using conditional quantile normalization (cqn v1.36.0) (Hansen et al., 2012) and the covariates sex, tissue of origin, genetic ancestry PC1 and PC2 and the first 15 technical sequencing PCS were regressed out. The network was constructed using the WGCNA package (v1.70.3) (Langfelder and Horvath, 2008) using a signed network for each timepoint separately. A modified version of robust WGCNA (rWGCNA) was used to reduce the influence of potential outlier samples on network architecture (Gandal et al., 2018). Samples were resampled from within each form of ASD ensuring that each iteration contained at least 4 samples from each form. In total 100 networks were constructed followed by consensus network analysis which takes the median of the topological overlap dissimilarity matrix across the 100 networks (Langfelder and Horvath, 2008). A soft-threshold power of 13,9,13 and 14 were used for days 25, 50, 75 and 100 respectively to achieve approximate scale-free topology (R^2^>0.8).

Modules were constructed using a minimal module size of 160, deep split = 4, cut height for creation of modules = 0.9999 and cut height for merging modules of 0.1. Each module eigengene (ME; first principal component of the module) was tested for association with the different forms of ASD as well as for association ASD overall using linear model. Module GO term enrichment was performed using clusterProfiler (v4.0.5) (Yu et al., 2012) with default parameters. Enrichment for common variation associated with ASD (Grove et al., 2019), SCZ (Pardinas et al., 2018), ADHD (Demontis et al., 2019), MDD (Howard et al., 2019) and BD (Mullins et al., 2021) was calculated using a stratified LDscore regression (v1.0.0) (Finucane et al., 2015). SNPs with 10 kb of a gene were assigned to that gene and the enrichment was calculated as the proportion of SNP heritability accounted for by the genes in the module divided by the proportion of total SNPs within the module. Cell type enrichment was performed using the bootstrap enrichment method from the EWCE package (v.1.0.1) (Skene and Grant, 2016) using control samples from previously published single cell data from human cortical spheroids set (Khan et al., 2020) with 100,000 permutations. As the background gene set, all genes expressed in both the current dataset and the single-cell dataset were used. Enrichment for high confidence ASD gene (gene score < 2 or syndromic genes) from the SFARI database (https://gene.sfari.org/database/gene-scoring/) and ASD postmortem modules (Parikshak et al., 2016) was performed using Fisher’s exact test. Module trajectories were modeled using polynomial splines with 2 degrees of freedom and 2 degrees of the piecewise polynomial and tested for significant differences using a linear model. Module preservation was performed using the modulePreservation function from the WGCNA package with 100 permutations.

Preservation was tested (1) at other timepoints within the current data set (2) in cortical samples from early developmental stages (8-16 weeks post conception) from the BrainSpan in vivo data set and (3) in other stem cell based neuronal models (Adhya et al., 2021; Flaherty et al., 2019; Lin et al., 2016; Schafer et al., 2019).

### Clustering stability

The different conditions (ASD form/Differentiation Day) were clustered based on the Spearman correlation of log_2_ fold-change of all expressed genes using the Ward.D2 clustering method on Euclidean distances as implemented in the ComplexHeatmap package (v2.9.3) (Gu et al., 2016). Cluster significance was tested using multiscale bootstrap resampling as implemented in the pvclust function in the pvclust package (v2.2.0) (Suzuki and Shimodaira, 2006) using the Ward.D2 clustering method on Euclidean distances with 10,000 bootstraps. The bootstrap probability measure is the frequency that a cluster appears in the bootstrap replicates.

To test for clustering similarity across clustering methods, clustering was performed using the Ward.D, Ward.D2, complete, average and Mcquitty methods on each of the Euclidean, maximum, Manhattan and Minkowski distances. For each pair of clustering and distance methods the cophenetic correlation was measured using the cor.dendlist function from the dendextend package (v1.14.0) (Galili, 2015). Cophenetic correlation is a measure of how well the pairwise distances of the unmodeled data is preserved in the dendrogram (Saracli et al., 2013). Additionally, the Rand Index was measured for each combination of these pairs using the rand.index function from the fossil package (v0.4.0) (Vavrek, 2011). Rand index is a measure of the similarity between two trials of clustering using the same data (Rand, 1971). To verify that the clusters were nor driven by a small subset of genes we subset the genes used for clustering to 10,000, 5,000 or 2,000 random genes, 100 times. The resulting genes were correlated using Spearman correlation and clustered using the Ward.D2 clustering method based on Euclidean distances. Cophenetic correlation and Rand indexes were calculated as above.

### Deconvolution

Bulk RNA sequencing counts were deconvolved using the control cell profiles from previously published single cell data (Khan et al., 2020). The reference-based decomposition was performed using the ReferenceBasedDecomposition function from the BisqueRNA package (v.1.0.5) (Jew et al., 2020). Changes in cell type proportion were tested using a logit mixed linear model. The logit transformation was performed using the logit function from the car package (v3.0.11) with a 0.001 adjustment. The model was implemented using the lme function from the nlme package (v3.1.152). Contrasts were used to compare each form of ASD at each time point to the control at the same time point using the glht function from the multcomp package (v1.4.18).

### Transcription factor binding over-representation

To identify transcription factor binding motifs (TFBS) over-represented in each module we used RcisTarget (v1.6.0) (Aibar et al., 2017). The transcription factor data base used was hg38 and included 10Kb upstream and downstream of the transcription start site (version 9; hg38 refseq-r80 10kb_up_and_down_tss.mc9nr). Only high confidence annotated TFs were used for downstream analysis. The regulatory relationship between modules was calculated as a kME (which is the correlation of each gene in the module with the module eigengene) weighted proportion of the number of genes regulated by the upstream module in the downstream module such that: 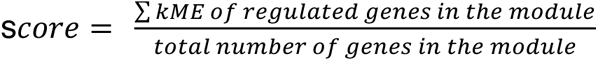. This weighted proportions puts larger weight on genes that are highly connected within the module. A module was considered to be highly regulated by another module if the kME weighted score was above 1 standard deviation above the mean (which corresponded to a weighted score of 0.341).

### Protein–protein interactions

Protein–protein interactions (PPI) were mapped using the Dapple (Rossin et al., 2011) module (v0.19) in GenePattern (genepattern.broadinstitute.org) using the hg19 genome build with 1000 permutations.

### Immunoprecipitation – Mass spectrometry

To validate predicted PPIs, we generated mass spectrometry data from key regulator proteins predicted to form a PPI. We tested antibodies against RELA (Active motif 40916), TP53 (Abcam ab26), CNOT6 (CST #13415), TBP (CST #8515), POLR2A (PTG 20655-1-AP), SOX9 (CST #82630), SMARCB1 (CST #91735), SMARCA4 (Abcam ab110641), and EP300 (Abcam ab275378). Antibodies that generated western blot bands at the appropriate size were selected for continued mass spectrometry (MS) analysis (Figure 6a). Mass spectrometry was then conducted for the 6 proteins (TBP, POLR2A, SOX9, SMARCB1, SMARCA4, and EP300) with western blot signal in NGN2-derived NPCs at day 4 (Nehme et al., 2018; Pintacuda et al., 2023; Pintacuda et al., 2021). MS experiments that yielded significance for the protein bait over IgG controls were then included in analyses (all proteins except SOX9).

Immunoprecipitations were done as in Pintacuda e al., *Cell Genom* 2023 (Pintacuda et al., 2023). Briefly, Neural Progenitor Cells (NPCs) were harvested at day 4 following NGN2 expression and dual SMAD/WNT inhibition in the presence of doxycycline (rtTA, tetON-NGN2). Cells were pelleted, washed with PBS and resuspended in 10x packed cell volume (PCV) IP lysis buffer (Thermo Scientific) with added Halt protease and phosphatase inhibitors (Thermo Scientific) for 20 minutes at 4°C. Cells were then centrifuged (16200g, 20min, 4°c) and resuspended in 3X PCV. Samples were quantified using the Thermo BCA protein assay.

For western blot analysis of the proteins of interest, cell lysates were diluted in 6X SMASH buffer ( 5mM Tis HCL ph 6.8, 10% glycerol, 2% SDS, 0.02% bromophenol blue, and 1% b-mercaptoethanol) and boiled for 10min at 95°c. Samples were then separated on a NuPAGE 4-12% Bis-Tris Protein Gel (Invitrogen) and transferred to a PVDF membrane (Life Technologies, 100V, 2 hours). Membranes were incubated at room temperature for one hour in TBS-T (0.1% Tween) with 5% BioRad Blotting-grade Blocker and then incubated overnight with antibodies listed above at 4°c. After 3x 10 min washes in TBST blots were incubated with horseradish peroxidase-conjugated secondary antibodies for 45min and again washed 3x (10min, TBST).

Western blots were imaged using SuperSignal West Femto Maximum Sensitivity Substrate (Thermo Scientific). Western blots were considered successful if the band was visualized at the expected molecular weight (Figure a). For the 7 proteins that showed successful western blots, 1-2 mg of protein extract was incubated with primary antibody (1-2 ug) at 4°c overnight. Following primary antibody incubation 100 ul of Protein A/G beads (Pierce) were added and incubated for 4 hours at 4°c followed by one wash with lysis buffer (Pierce) with added Halt protease and phosphastase inhibitors (Thermo Scientific) and two washes with PBS and resuspended in PBS for Mass spectrometry.

### Mass Spectrometry and Quantification

Samples were processed and quantified for Mass Spectrometry as previously described (Pintacuda, Cell Genomics 2023). Briefly, immunoprecipitations were eluted using SDS and subject to electrophoresis for gel extraction. Bands were excised and cut into approximately 1 mm^3^ pieces. Gel pieces were then subjected to a modified in-gel trypsin digestion procedure, washed and dehydrated with acetonitrile for 10 min, followed by removal of acetonitrile. Pieces were then completely dried in a speed-vac. Rehydration of the gel pieces was with 50 mM ammonium bicarbonate solution containing 12.5 ng/*µ*l modified sequencing-grade trypsin (Promega, Madison, WI) at 4°C. After 45 minutes, the excess trypsin solution was removed and replaced with 50 mM ammonium bicarbonate solution to just cover the gel pieces. Samples were then placed in a 37°C room overnight. Peptides were later extracted by removing the ammonium bicarbonate solution, followed by one wash with a solution containing 50% acetonitrile and 1% formic acid. The extracts were then dried in a speed-vacuum (∼1 hr). The samples were then stored at 4°C until analysis.

On the day of analysis, the samples were reconstituted in 5 - 10 *µ*l of HPLC solvent A (2.5% acetonitrile, 0.1% formic acid). A nano-scale reverse-phase HPLC capillary column was created by packing 2.6 *µ*m C18 spherical silica beads into a fused silica capillary (100 *µ*m inner diameter x ∼30 cm length) with a flame-drawn tip. After equilibrating the column each sample was loaded via a Famos auto sampler (LC Packings, San Francisco CA) onto the column. A gradient was formed, and peptides were eluted with increasing concentrations of solvent B (97.5% acetonitrile, 0.1% formic acid). As peptides eluted, they were subjected to electrospray ionization and then entered an LTQ Orbitrap Velos Pro ion-trap mass spectrometer (Thermo Fisher Scientific, Waltham, MA). Peptides were detected, isolated, and fragmented to produce a tandem mass spectrum of specific fragment ions for each peptide.

Peptide sequences (and hence protein identity) were determined by matching the UniProt human protein database (release 2023_01) with the acquired fragmentation pattern using Sequest (Thermo Fisher Scientific, Waltham, MA) (Eng et al., 1994). The database included a reversed version of all the sequences and the data was filtered to between 1-2% peptide false discovery rate. Protein quantification was performed using GFY Core Version 3.8 (Harvard University, Cambridge, MA). IP-MS data analysis was described in (Pintacuda et al., 2023).

### Protein-Protein Interaction Analyses

To determine if the M5 proteins formed a significant interaction network, we compared the logFCs between each ‘bait’ and the other interactors predicted to be in the M5 network, and found as a protein in any of the MS experiments (943 unique proteins identified across all IP-MS experiments). If the interactors were not pulled down by the bait, the logFC was considered 0. We used the mean logFC across baits as a measure of average connectedness of the network.

To assess the statistical significance of these networks, we ran 10,000 permutations by randomly selecting proteins of the same size as the network from the remaining set of 943 uniquely identified proteins. We compared the average logFC of each of these randomly generated networks to create a population curve. We then examined the z-scored connectedness of the network. Gene-lists used to compare networks were SFARI genes (levels 1,2,S), ASD risk genes (Ruzzo et al., 2019a; Satterstrom et al., 2020), ID risk genes (Leblond et al., 2021), NDD risk genes (Leblond et al., 2021) and DD-enriched, ASD-enriched, and NSD-nonspecific (Fu et al., 2022).

### gRNA Design and Crop Seq cloning

Three gRNAs/target as well as 39 total Non-Targeting Controls (NTC) were designed using the CRISPick database (https://portals.broadinstitute.org/gppx/crispick/public). Sequences were designed as follows: A 5’ overhang (ATCTTGTGGAAAGGACGAAACACC) and a 3’ overhang (GTTTAAGAGCTATGCTGGAAACAGCATAGCAAGT) that is compatible with the CROP-seq vector were added on the outside of the gRNA sequence. A “G” nucleotide was also incorporated between the 5’ overhang and the gRNA sequence. All oligos were ordered as a pool (Integrated DNA Technologies). The CROP-seq vector backbone (Datlinger et al., 2017) was obtained from addgene (addgene: #86708) and digested using New England Biolabs’ (NEB) BsmBI enzyme. The oligo pool was diluted at a concentration of 100uM, PCR amplified, (Q5® Hot Start High-Fidelity 2X Master Mix) gel extracted, and cleaned. The vector and the pooled oligo insert was then assembled (NEBuilder HiFi DNA Assembly Reaction). To validate the insertion was present post assembly, a PCR amplification using the U6 outer forward primer (TTTCCCATGATTCCTTCATATTTGC) and gRNA end reverse primer (AGTACAAGCAAAAAGCAGTGTCTCAA) was run and visualized. The pooled vectors were then transformed into NEB Stable Competent E.Coli (High Efficiency), and plated at various dilutions on LB - Ampicillin 1:1000 plates overnight at 30°C. Colonies were picked and PCR amplification of the insert was done to insure its presence. All bacterial colonies were then scraped from the plate and grown using 200ml of 2.5% LB broth + 1:1000 Ampicillin. The broth containing colonies is then shaken at 170 RPM at 30°C and completed once the OD reaches 2.0. DNA was then isolated via the Qiagen Maxi Plasmid Kit.

### Lentiviral Production

HEK293T cells were expanded in DMEM, 10% FBS, and 1% Pen-Strep (HEK medium). 5×10^6^ HEK293T are plated onto two poly-ornithine (5ug/ml) coated 10 cm plates for 48 hours. Media from both plates was replaced with opti-mem before cells from one 10cm plate were transduced with a lentiviral vector (pLV-KRAB-dCas9 (addgene #71236) or CROP-dsRed (addgene #201999)) while the other plate of cells was transfected with 15ug of the pooled plasmid library, 15ug of psPAX2, and 1.5 ug of VSV-G by way of the lipofectamine 3000. 16 hours later, the Opti-mem media was replaced with HEK medium and that medium was collected and replaced every 24 hours for two days. The preserved HEK medium was then spun down with 12.5 ml of lenti-X concentrator (Takara Bio), in an effort to capture the two distinct lentiviruses, at 1500g for 45 minutes. The supernatant was discarded and the pellet was resuspended in 400ul of PBS, aliquoted, and stored at −80°C.

### Derivation of NGN2-induced neural cells, transduction and FACS

We derived NGN2-induced hNPCs according to (Guss et al., 2024). A 6 well plate was coated with Matrigel (1:50 in Knockout DMEM) for 2 hours at 37°C. A dox-inducible NGN2 iPSC line, WTC11, was seeded at 75,000 cells/cm^2^ in MTeSR+ supplemented with 10uM of Y-27632. 24 hours afterwards, cells were subjected to Induction Medium (1:100 Glutamax, 1:66.7 20% Glucose in DMEM:F12, N2 supplement 1:100, Doxycycline 2ug/ml, LDN-193189 200nM, SB431542 10uM, XAV939 2*µ*M). Day 2 Involves the induction and selection using the following media: 1:100 Glutamax, 1:66.7 20% Glucose in DMEM:F12, N2 supplement 1:100, Doxycycline 2ug/ml, LDN-193189 100nM, SB431542 5uM, XAV939 1*µ*M, and Puromycin 5ug/ml. 24 hours post selection, the media is changed fully with NES Complete & selection media (DMEM:F12, Glutamax 1:50, Pen.Strep 1:100, MEM NEAA 1:100, B27 w/o Vitamin A 1:50, N2 Supplement 1:100, EGF 10ng.ml, bFGF 10ng/mL, puromycin 5ug/ml, and Y-27632 1:1,000. One day later and for the duration of the culture, cells are put on NES complete medium which is the same as listed above apart from the use of Y-27632 and puromycin.Once hNPCs have been generated, 500k hNPCs are seeded in every well of a 6 well plate with NPC maintenance media (n= 6 replicates). To each well, GFP-KRAB-dCas9 (40 uL/well of a 6 well) plasmid and CROP-dsRed-gRNA (33 uL/well of a 6 well) were added. Media was replaced daily for 7 days before cells were collected for FACs. For FACS, hNPCs were washed with PBS, dissociated into single cell suspension using Accutase, and resuspended in 500ul FACS buffer per replicate (7.5% BSA, .5 M EDTA, RNase inhibitor 1:200, DAPI .5mg/ml). A population of NPCs that were not transfected with plasmid were used as a negative control to establish FACs gating. Double positive GFP/RFP cells were sorted for downstream single cell analyses (∼30,000 cells per replicate).

### Single cell RNA seq

Sorted cells then underwent single cell RNA extraction using the 10x genomics Chromium Next GEM Single Cell 3’ GEM, Library & Gel Bead Kit v3.1(PN-1000121). The 10x protocol for library prep was adapted from Gasperini et al., 2019 (Gasperini et al., 2019). gRNA libraries were sequenced separately from scRNA (GEX) libraries. ScRNA and gRNA libraries were aligned to the human genome (GRCh38) with CellRanger (7.0.1). Bam files generated from gRNA libraries were then analyzed as in (Hill et al., 2018) Github: https://github.com/shendurelab/single-cell-ko-screens). In short, a text file of gRNA sequences is supplied as well as the common sequence appearing prior to the gRNA sequence and each of these reads is counted from the bam file and paired to the cells within the scRNA libraries via cell UMI matching. Only cells containing gRNA barcodes with a UMI count of at least 10 were kept in the downstream analysis. This cut-off has also been previously used (Hill et al., 2018) and was determined to be sufficient for significant decrease in expression of the target. All analysis was performed on cells that expressed the gRNA of a single target. Single cell data was then processed using standard best practices for Seurat v4 including DoubletFinder to remove doublets, log-normalization, and integration. Normalization using SCTransform was followed by clustering. Canonical markers were used to determine cell-type classification including “NPC” markers SOX2, MKI67, VIM, and HES1 and cycling neural progenitor cells (cNPCs) expressing G2 mitotic markers: TOP2A, HMGB2, CDK1, NUSAP1, UBE2C, and BIRC5. A standard pseudobulk differential expression was performed using the libra package function, run_de(). The method used was edgeR and the type was QLF.

---

**Extended Data Figure 1. Experimental design.**

(a) Schematic representation of hCO differentiations from each derived hiPS cell line from each individual.

(b) Schema of dropped lines due to quality control metrics

**Extended Data Figure 2. Sequencing depth in CNV loci**.

Whole genome sequencing depth in (a) 16p11.2 (b) 15q13.3 (c) 22q11.2 and (d) 22q13.3 loci averaged over 10kb windows.

**Extended Data Figure 3. Point mutations identified using whole genome sequencing.**

(a) Point mutations in CACNA1C (G406R) identified in the 3 lines from individuals with Timothy syndrome. (b) Point mutations in PCDH19 (Y243* and Q201P) identified in the 2 lines from individuals with PCDH19 syndrome. (c) Point mutations in SHANK3 (R522W) identified in the 2 lines from the individual with SHANK3 mutation. All mutations were visualized with the Integrative Genomics Viewer (IGV).

**Extended Data Figure 4. Sources of variation in the gene expression data**

(a) Hierarchical clustering of conditions (ASD form and differentiation day) based on the median gene expression of all samples in each condition showing that the samples cluster by differentiation day and the genetic form of ASD. Top color bar denotes ASD form and lower bar denotes differentiation day.

(b) Association of the top 20 gene expression principal components (PC) with the different covariates. Numbers in brackets on the y-axis are the percent of variance explained by these PCs. Differentiation time is highly associated with the 1^st^ PC (Adjusted R^2^ = 0.92).

(c) Variance explained by each of the covariates for each gene. Distributions represent the density of the percent of variance explained. The median value of percent of explained variance for each variable are denoted below the plots.

Boxplots in d and e show: center – median, lower hinge – 25% quantile, upper hinge – 75% quantile, lower whisker – smallest observation greater than or equal to lower hinge –1.5× interquartile range, upper whisker – largest observation less than or equal to upper hinge +1.5× interquartile range.

**Extended Data Figure 5. Log fold change in the affected CNVs and genes carrying point mutations.**

Genes inside the CNV locus or genes carrying the point mutations and marked in blue. Genes significantly differently expressed over all time points are denoted by asterisks. The percent of genes in the CNV which were DE across the 4 timepoints was as follows: 15q13 - 81.8% of genes, 16p11del - 61.5% of genes, 16p11dup - 50% of genes, 22q11del - 88.9% of genes, 22q13del - 77.3% of genes.

**Extended Data Figure 6. Validation of clusters based on correlation of logFC.**

(a) Cophenetic correlation and Rand index across multiple clustering and distance methods (left) and gene subsampling (right). Cophenetic correlation measures how faithfully the pairwise distances of the unmodeled data are preserved in the dendrogram. Rand index measures the similarity between two different clusterings of the same data.

(b) Cluster significance using multiscale bootstrap resampling. P values are indicated in red for each branching point. Bootstrap probability measures the frequency that a cluster appears in bootstrap replicates.

**Extended Data Figure 7. Annotation of differentially expressed genes.**

(a) Overlaps between differentially expressed genes at all time points and risk genes for ASD from either SFARI or from large scale whole exome (WES) sequencing studies (Ruzzo et al., 2019a; Satterstrom et al., 2020) as well as with neurodevelopmental disorders (NDDs) and intellectual disability (ID) risk genes (Leblond et al., 2021). Color represents the OR and the size of the point represented the −log_10_ FDR. Only positive significant overlaps (OR > 1 and FDR < 0.05) are shown.

(b) Select gene ontology (GO) terms enriched in upregulated (red) or downregulated (blue) genes in each of the ASD forms at day 25. Color corresponds to normalized enrichment score (NES). Point size corresponds to the level of significance (-log_10_(FDR)).

* FDR < 0.05, ** FDR < 0.01, *** FDR < 0.005.

**Extended Data Figure 8. Single cell analysis of human cortical organoid control cells from Khan et al. (Khan et al., 2020).**

(a) Clusters identified using samples from Khan et al 2020. Clustering was based on all samples while cluster characterization was based on controls samples only. Selected markers for (b-c) Cycling progenitor (CycPro), (d-e) Radial glia (RG-1, RG-2, and RG-3), (f-g) outer radial glia (oRG), (h-i) intermediate progenitors (IPC), (j-k) Excitatory neurons (ExNeu-1 and ExNeu-2), (l-m) Intermediate neurons (InNeu)

**Extended Data Figure 9. Deconvoluted cell proportions.**

Cell proportion in the different forms of ASD at different time points during hCO differentiation. Significant changes are marked by asterisk. CycPro - cycling progenitor, RG-1 - radial glia cluster 1 (early), RG-2 - radial glia cluster 2 (late), RG-3 – radial glia cluster 3, IPC - intermediate progenitor cells, ORG - outer radial glia, ExNeu-1 - excitatory neurons cluster 1 (early), ExNeu-2 - excitatory neurons cluster 2 (late), InNeu - Intermediate neurons. * - FDR < 0.05.

**Extended Data Figure 10. Association of ICs with ASD form and differentiation day.**

Hierarchal clustering of the association of independent components (IC) with the different forms of ASD at all time points. * - FDR < 0.05.

**Extended Data Figure 11. Construction of WGCNA modules**

Dendrogram from which the WGCNA modules at day 25 were derived using the dynamic tree cut algorithm (Langfelder et al., 2008). Genes associations (adjusted R2) with covariates is also indicated.

**Extended Data Figure 12. Module preservation.**

Preservation of day 25 modules at other timepoints (left), in vivo (8-16 weeks post conception) (Li et al., 2018) and in other stem cell derived neuronal data sets: Adhya2020 (Adhya et al., 2021), Flahrety2019 (Flaherty et al., 2019), Lin2016 (Lin et al., 2016) and Schafer 2019 (Schafer et al., 2019). The Z_summary_ preservation score is indicated.

**Extended Data Figure 13. Module eigengene trajectories.**

(a) Module eigengene trajectories of day 25 modules across differentiation. Significance was tested using a linear model with polynomial splines with 2 degrees of freedom and 2 degrees of the piecewise polynomial.

(b) Module eigengene trajectories of the regulators in M5 (blue) and the ASD risk genes that are regulated by these regulators (grey).

* FDR < 0.05, ** FDR < 0.01, *** FDR < 0.005.

**Extended Data Figure 14. Immunoprecipitation followed by mass spectrometry (IP-MS) of M5 regulators.**

a) Volcano plots show logFC of proteins compared to IGG controls. Highlighted are SFARI genes (1,2,S).

b) Significance of connectedness of networks is determined by comparing the network mean logFC to 10000 randomly selected networks of the same size across each of the protein baits. Blue dotted line shows the network mean logFC and black line shows the 95^th^ percentile.

c) Connectivity networks from gene lists of interest. Left: Full connectivity of significant interactions of all baits. Size of circle represents number of significant proteins for each bait. Right: NDD and ASD risk genes (Leblond et al., 2021; Ruzzo et al., 2019b; Satterstrom et al., 2020)

**Extended Data Figure 15. M5 Regulator CRISPRi methods and validation.**

(a) Schema of CROP-Seq vector

(b) cells transduced by number of gRNA (x-axis). Only cells expressing gRNA for a single gene-target are retained for downstream analyses.

(c) Proportion of cells uniquely expressing guide RNAs for each target within each experiment (n=6 technical replicates, 3 guide RNAs per target). SMARCB1 and TP53 show increased proportion compared to non-targeting controls (NTCs). *p<0.05, **p<0.01.

(d) Single cell UMAPs of markers used to differentiate cycling and non-cyling NPCs

**Extended Data Figure 16. M5 Regulator CRISPi additional data**

(a) Broad positive correlation between DEGs across targets in NPCs (right) and cNPCs (left) and DEGs in patient-derived organoids across developmental stages.

(b) Enrichment of rare genetic variation gene-lists in the differentially expressed genes of each target

(c) Network plot of differentially expressed genes by hCO-derived WGCNA modules

**Extended Data Figure 17. Gene ontology of CRISPRi targets**

Gene Ontology for targets with >40 DEGs shows changes to synapse-related terms and cell fate, particularly in SOX9, RBBP5, and SMARCB1.

