## Extended Data Figures for "Developmental convergence and divergence in human stem cell models of autism spectrum disorder"

Extended Data Figure 1

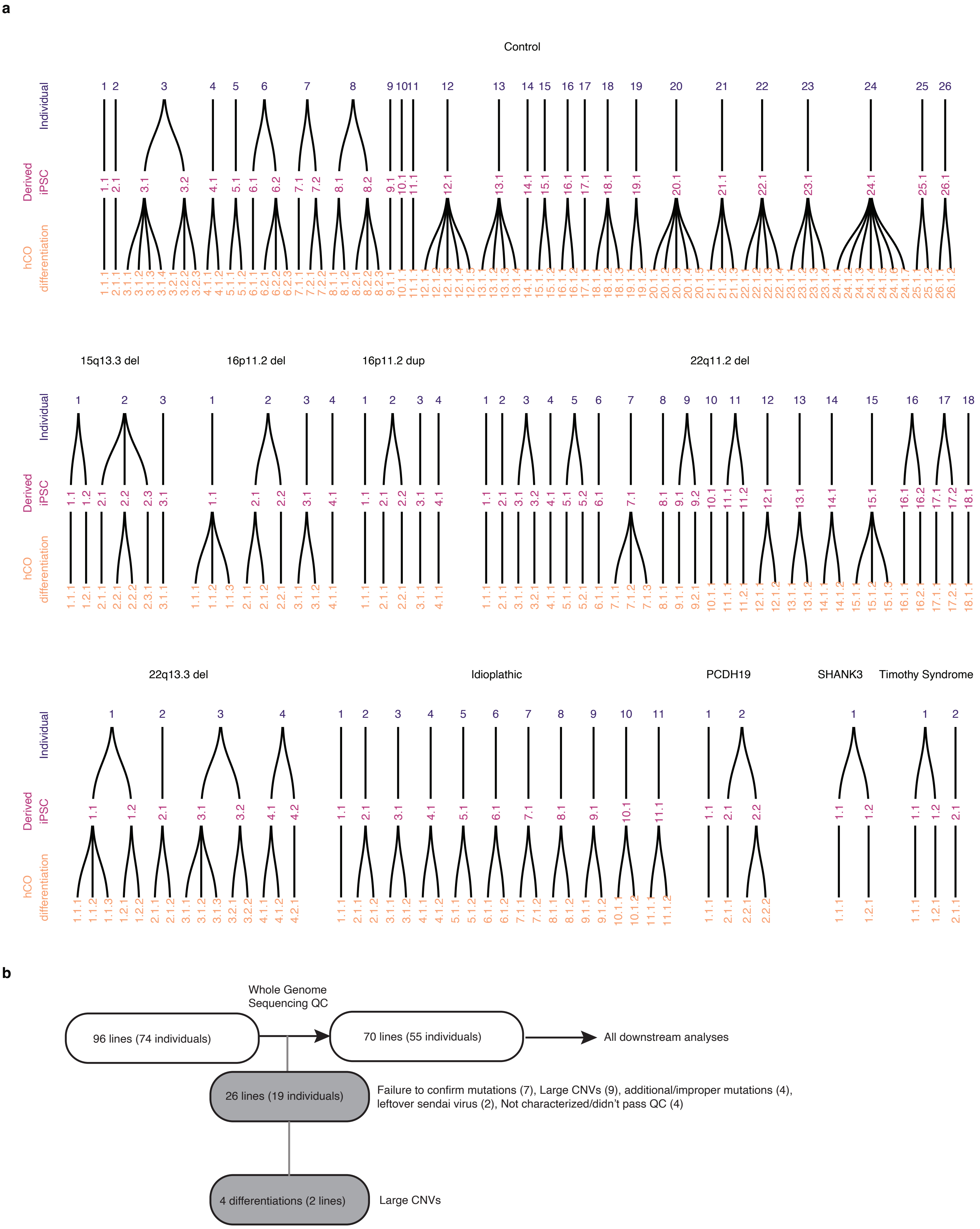

Extended Data Figure 2

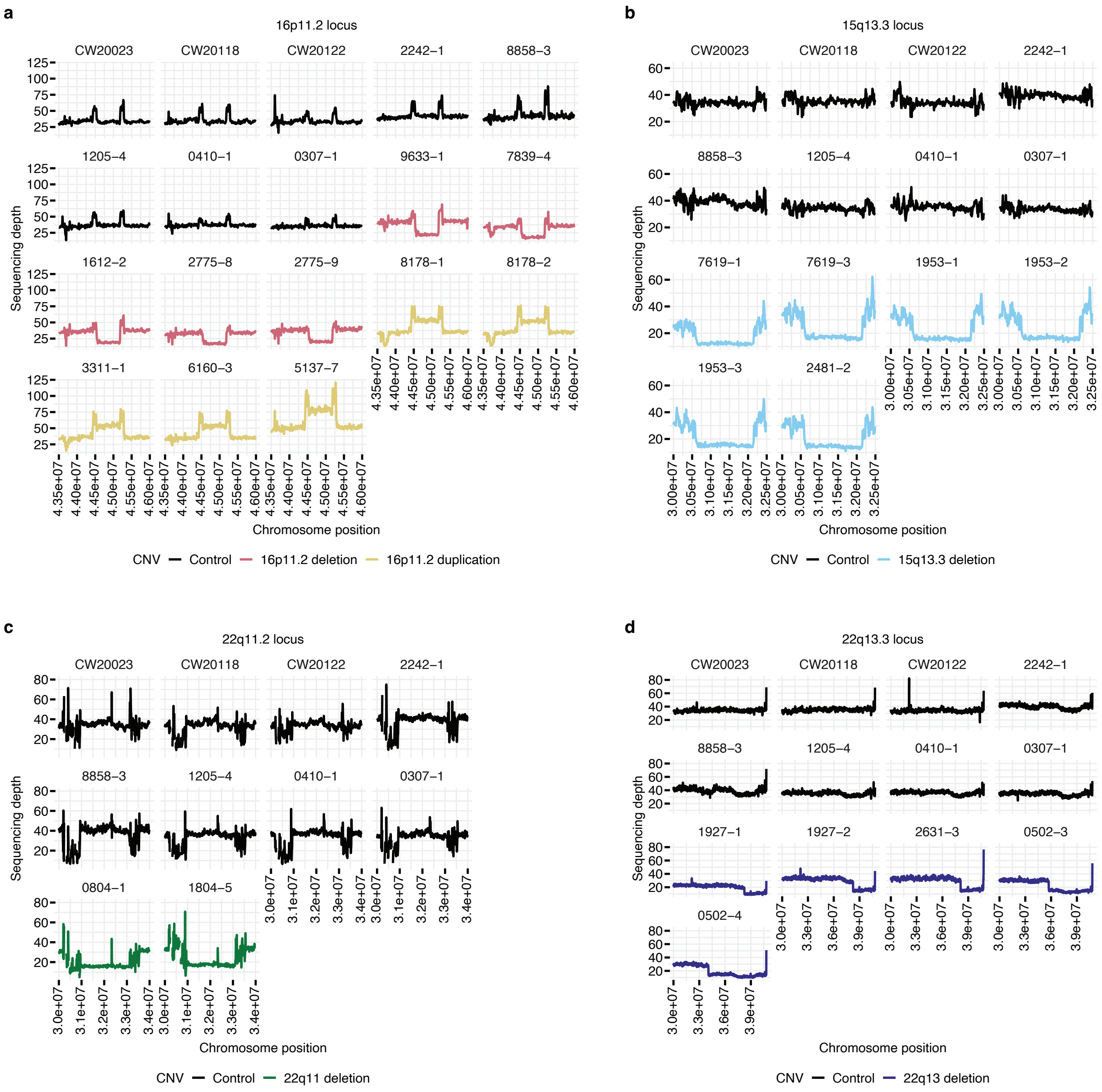

Extended Data Figure 3

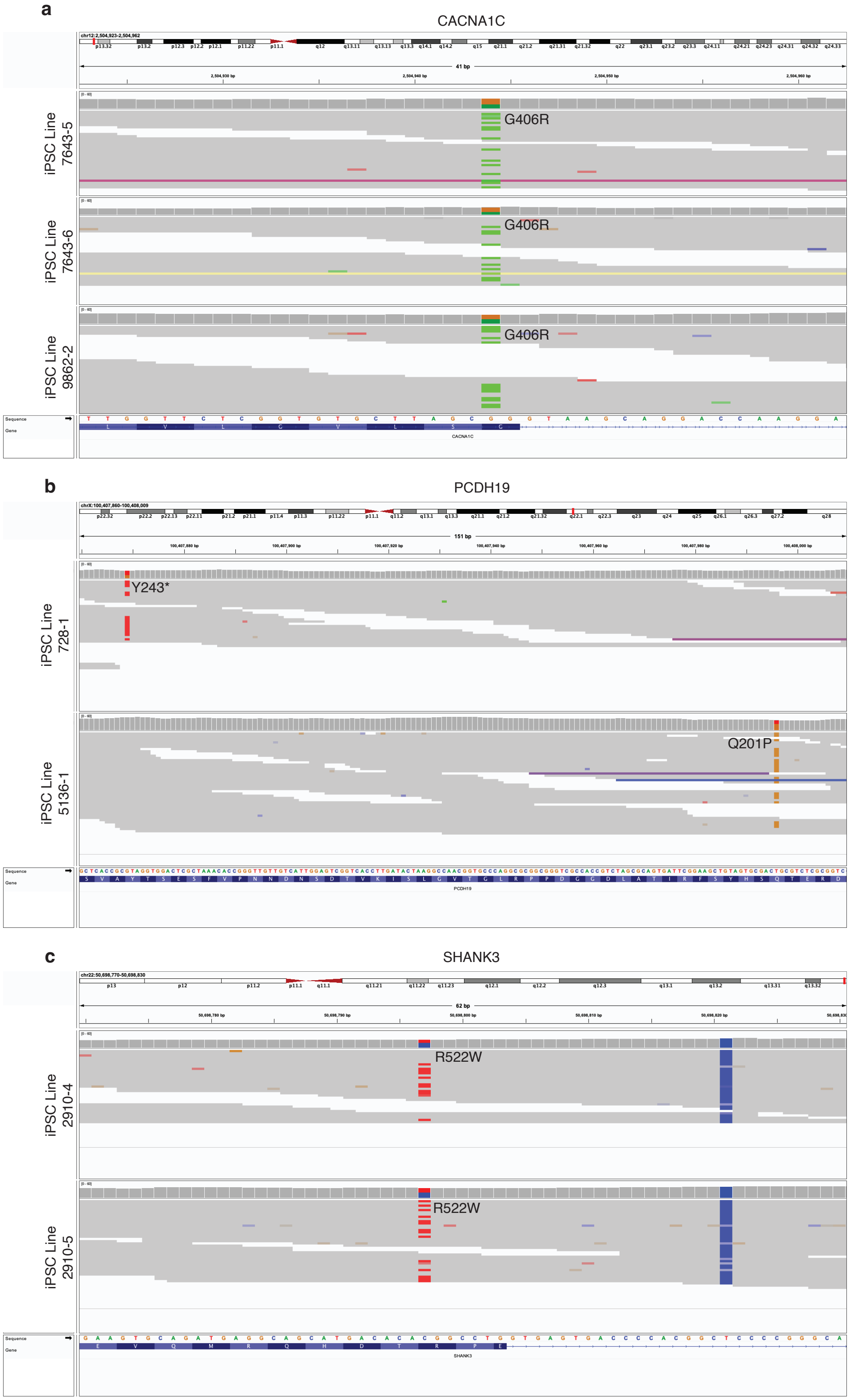

Extended Data Figure 4

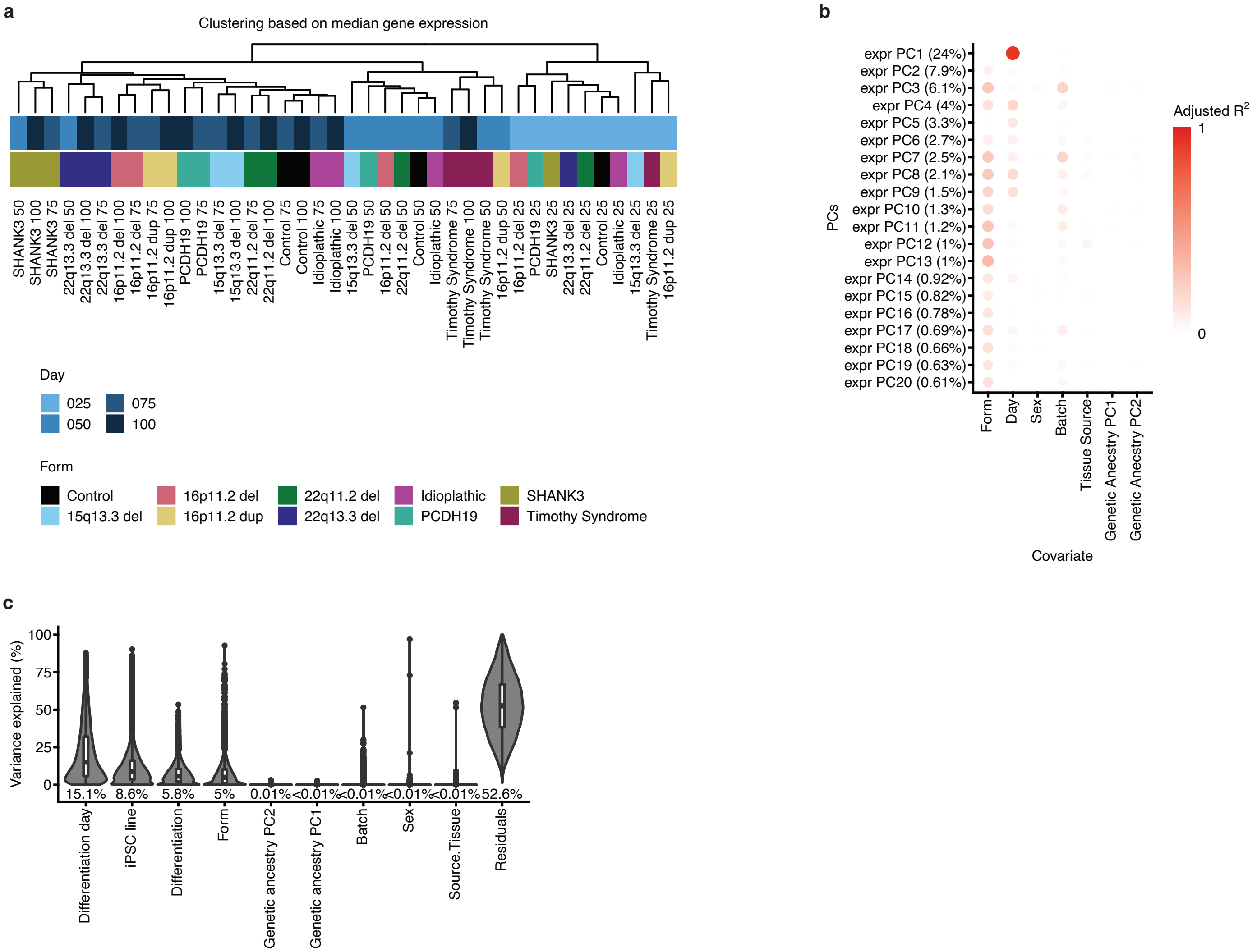

Extended Data Figure 5

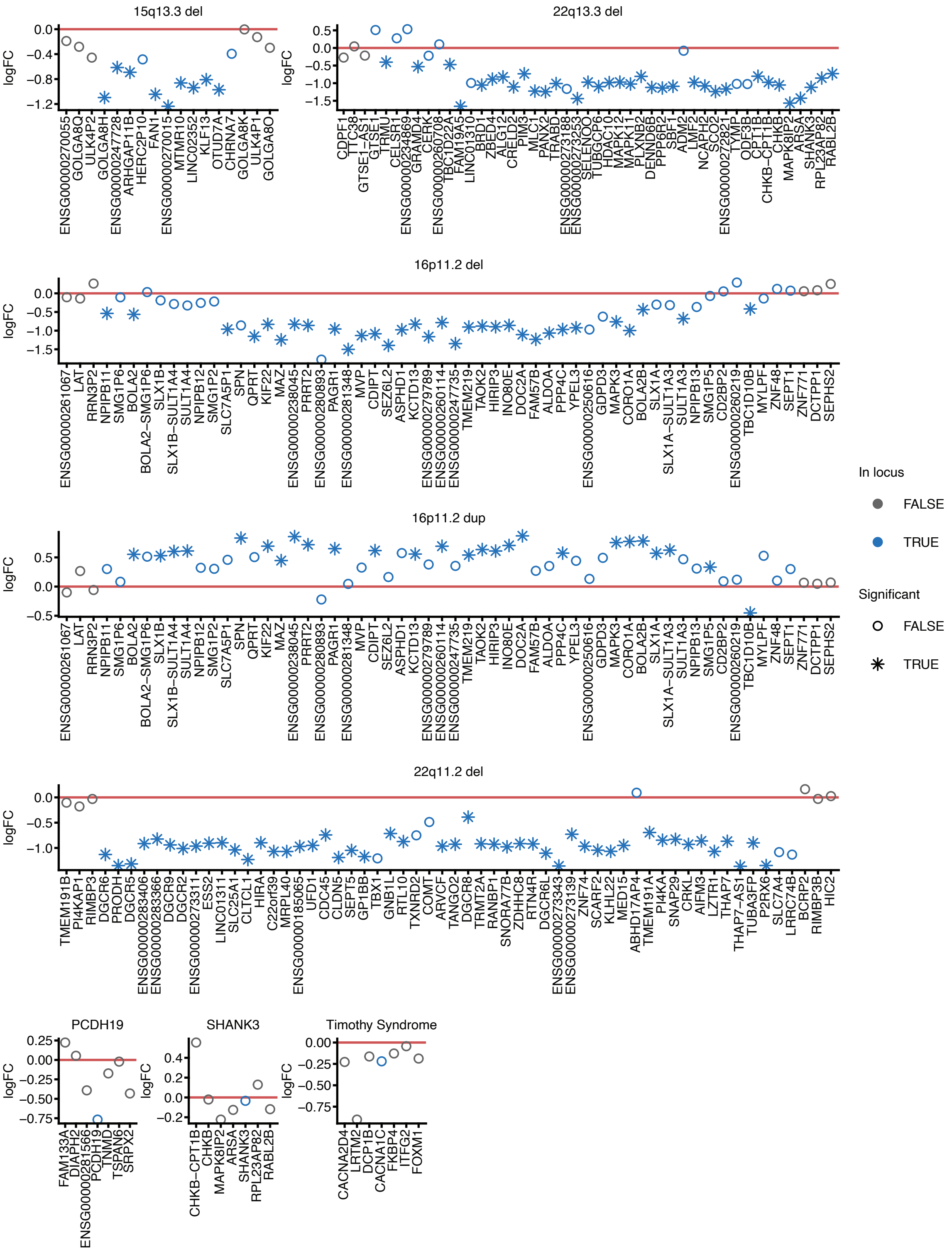

Extended Data Figure 6

**a**

Clustering simiarity across different clustering methods and gene subsets

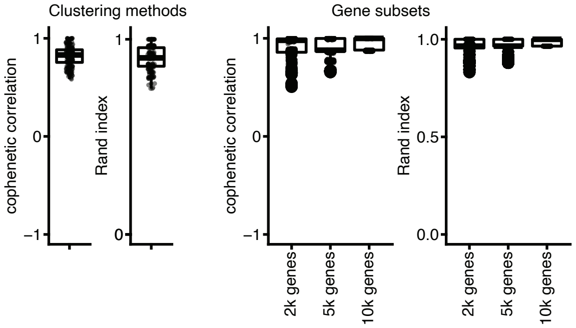


**b**

Edge P values using multiscale bootstrap resampling

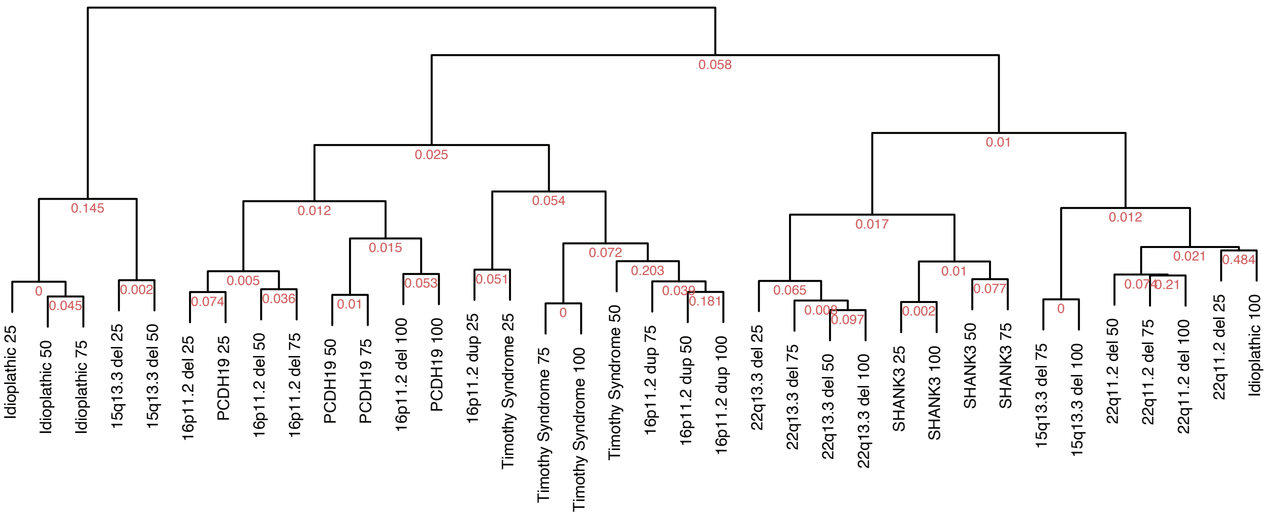

Extended Data Figure 7

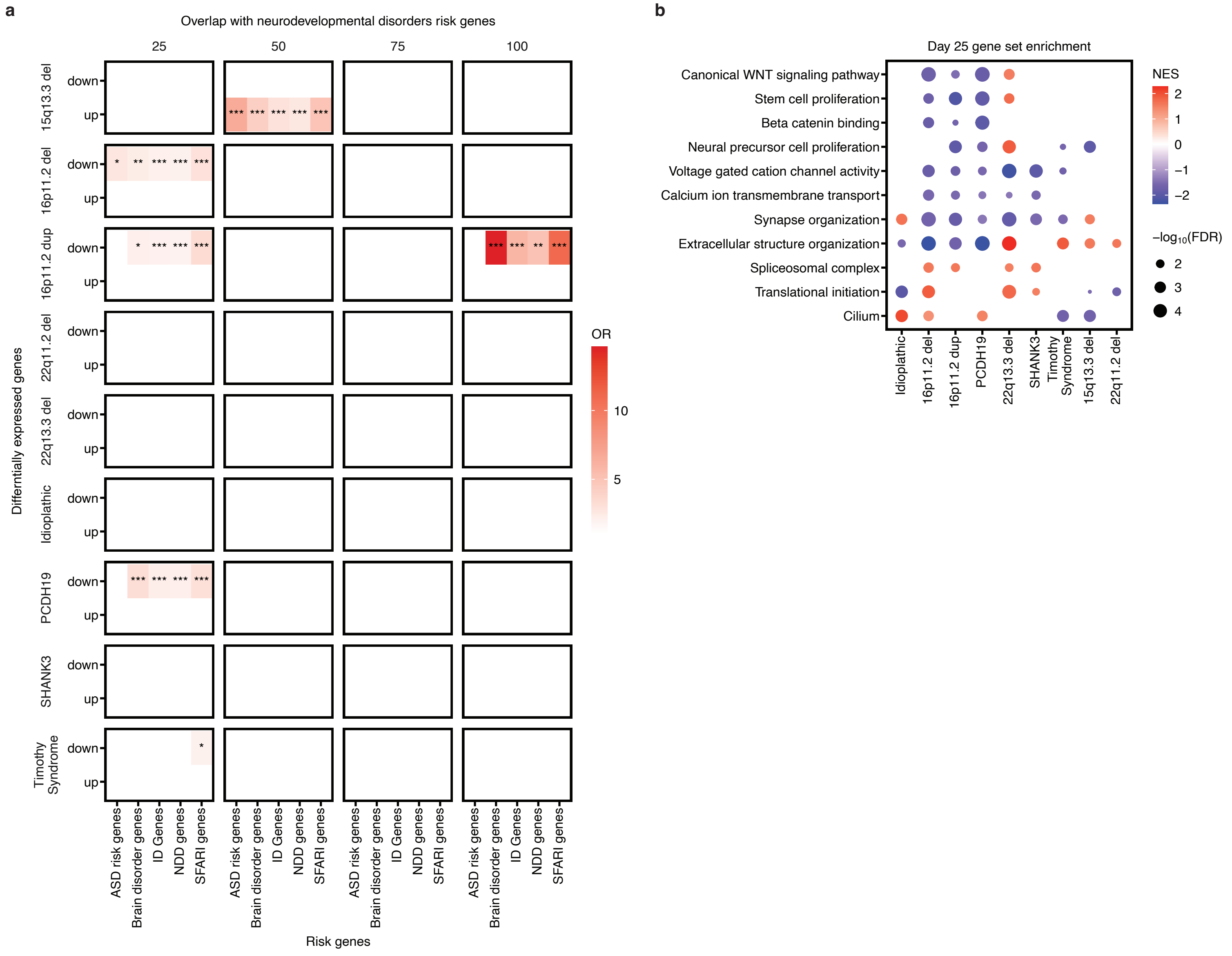

Extended Data Figure 8

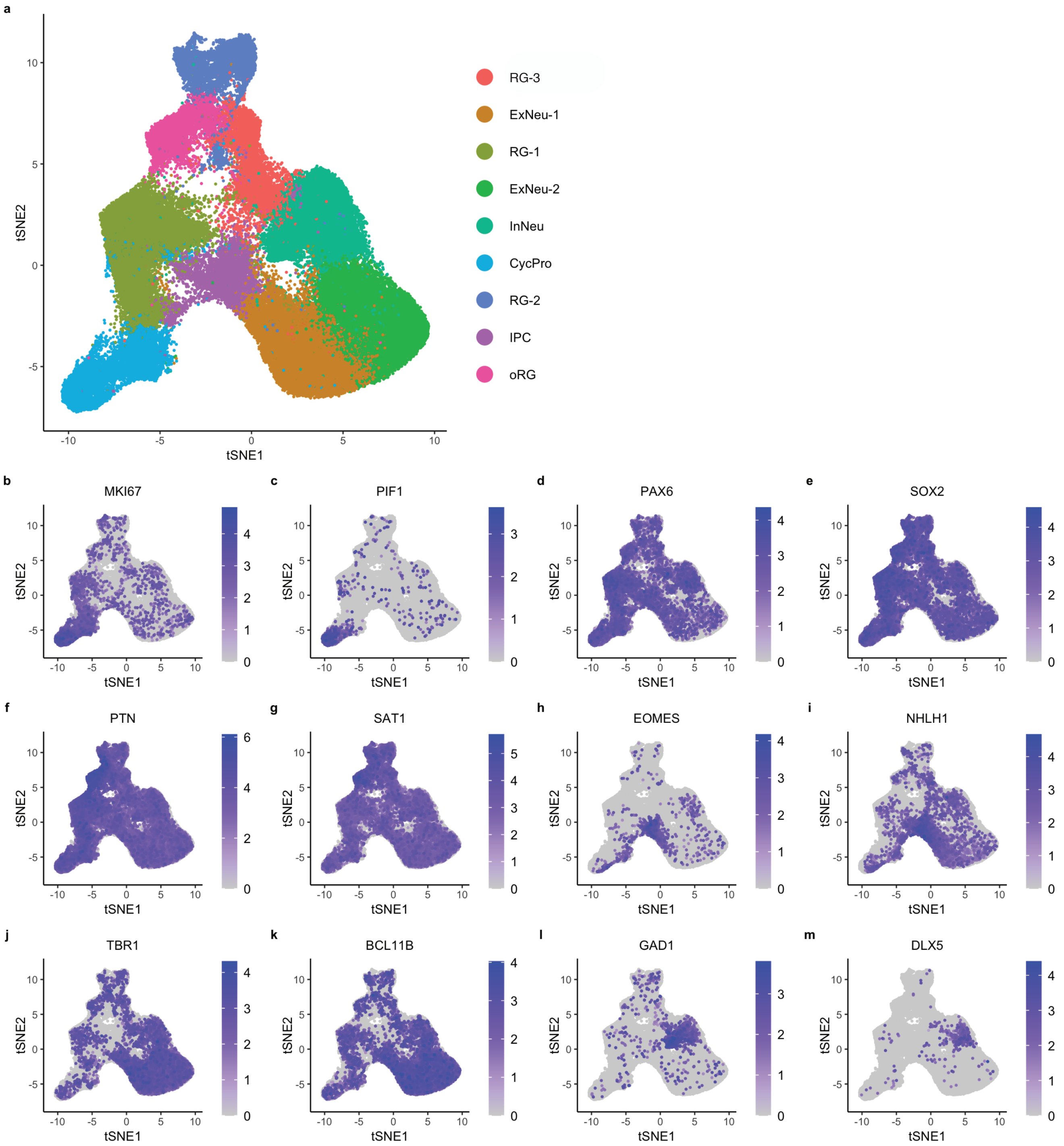

Extended Data Figure 9

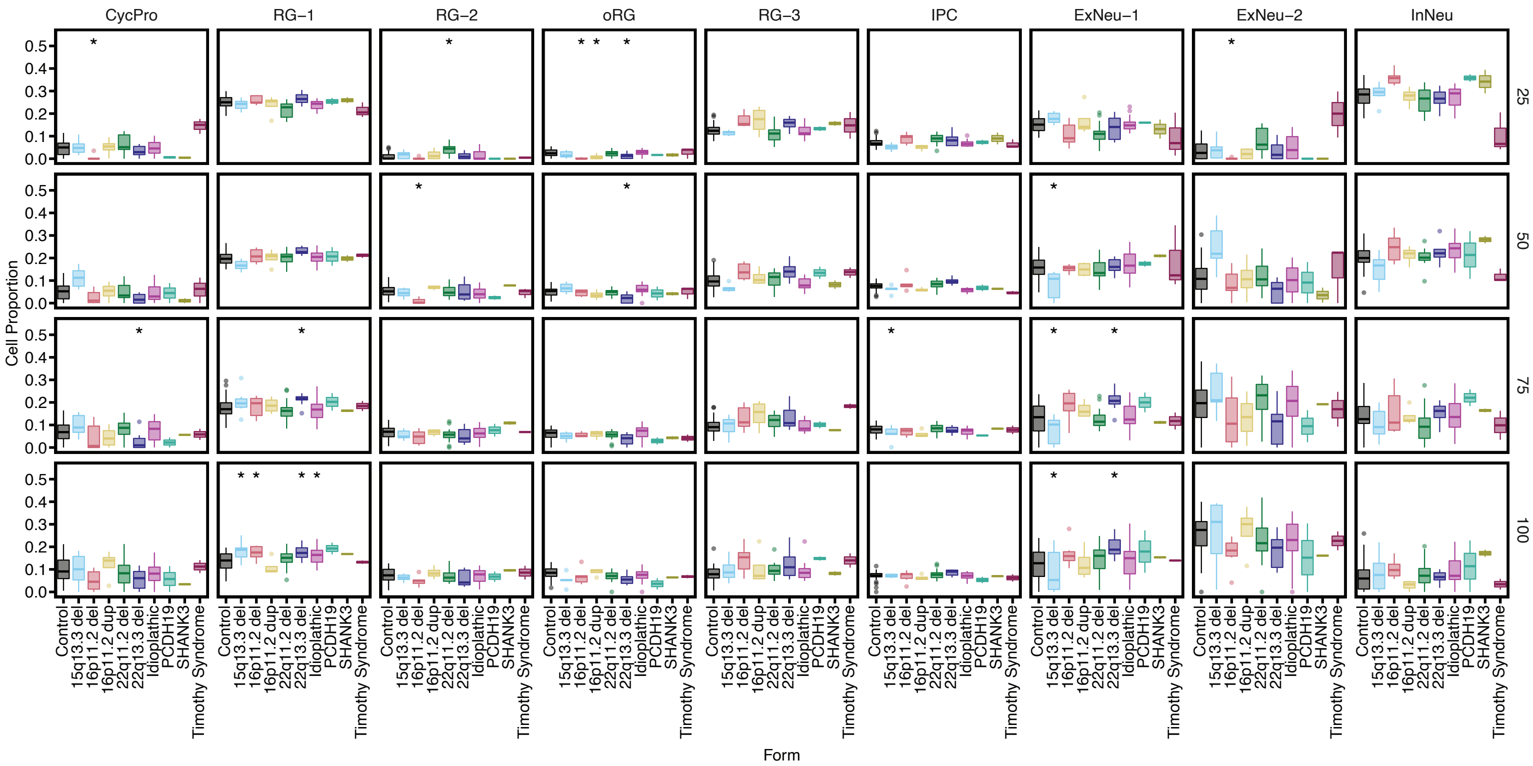

Extended Data Figure 10

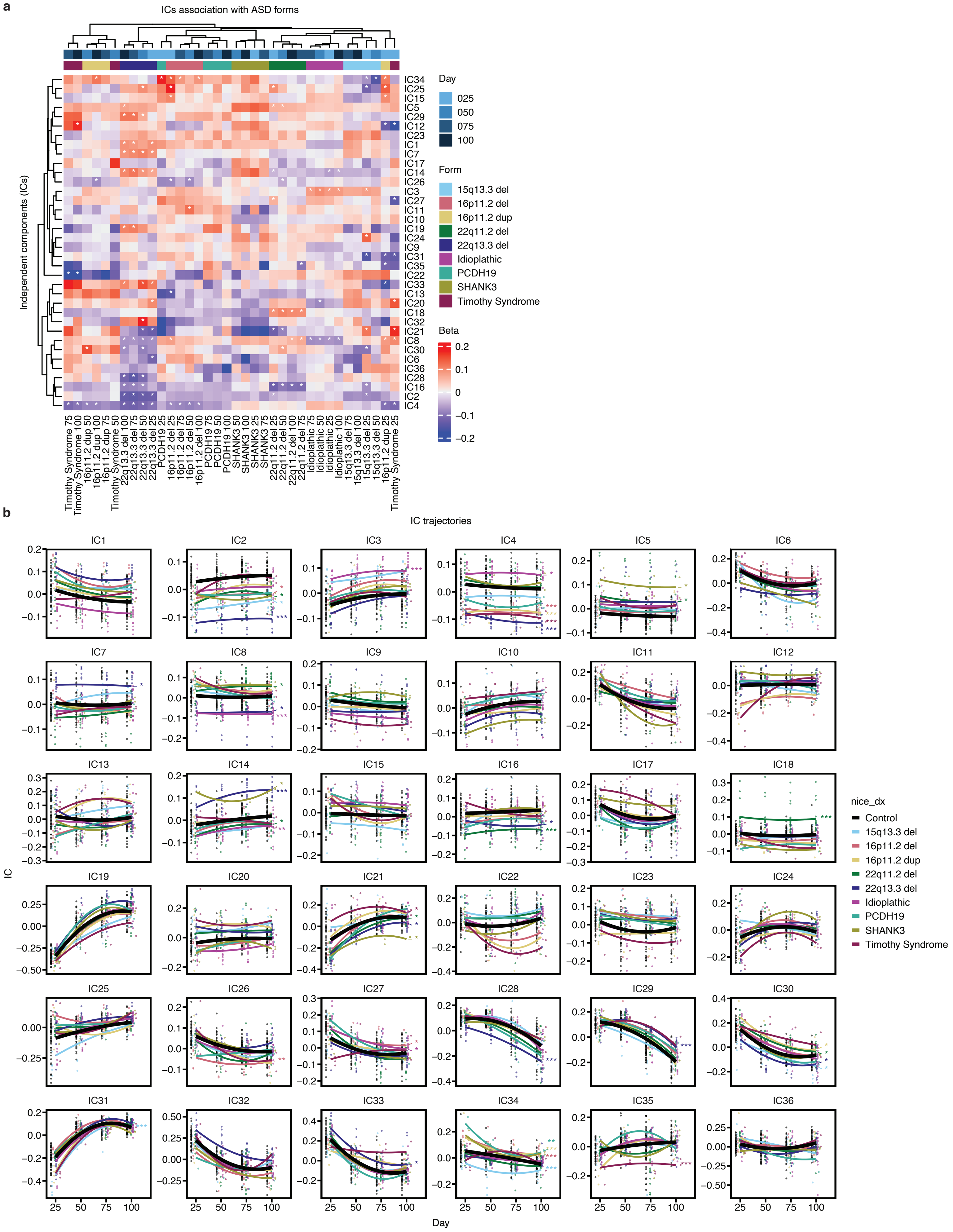

Extended Data Figure 11

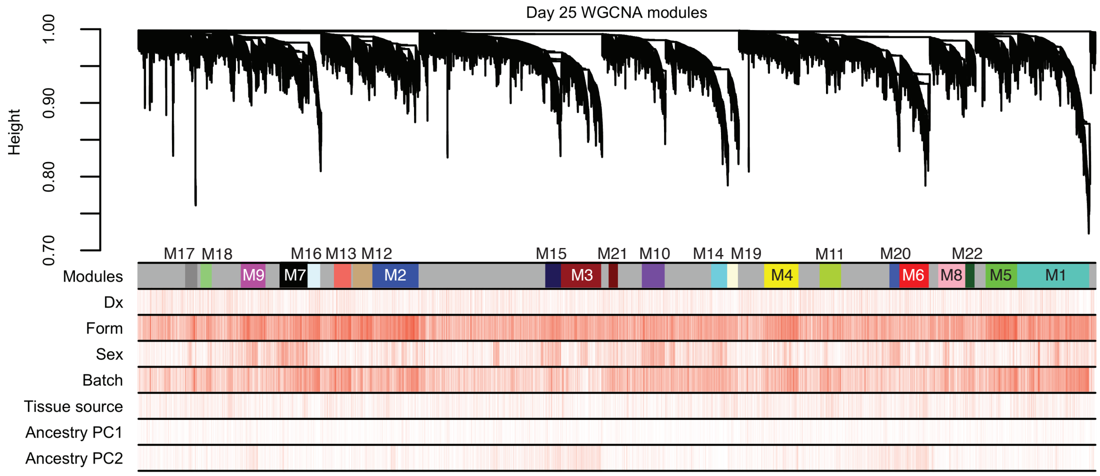

Extended Data Figure 12

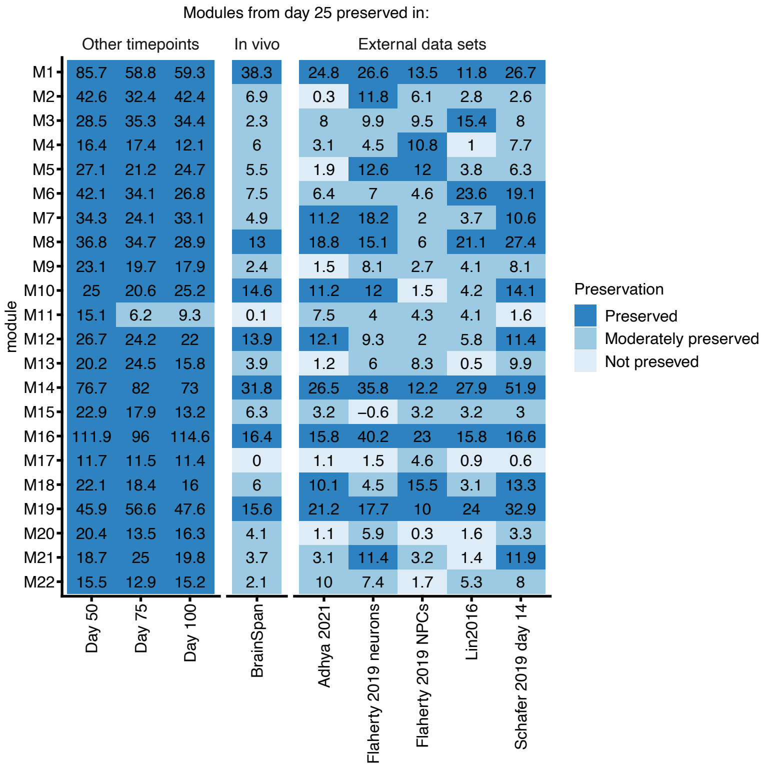

Extended Data Figure 13

**a**

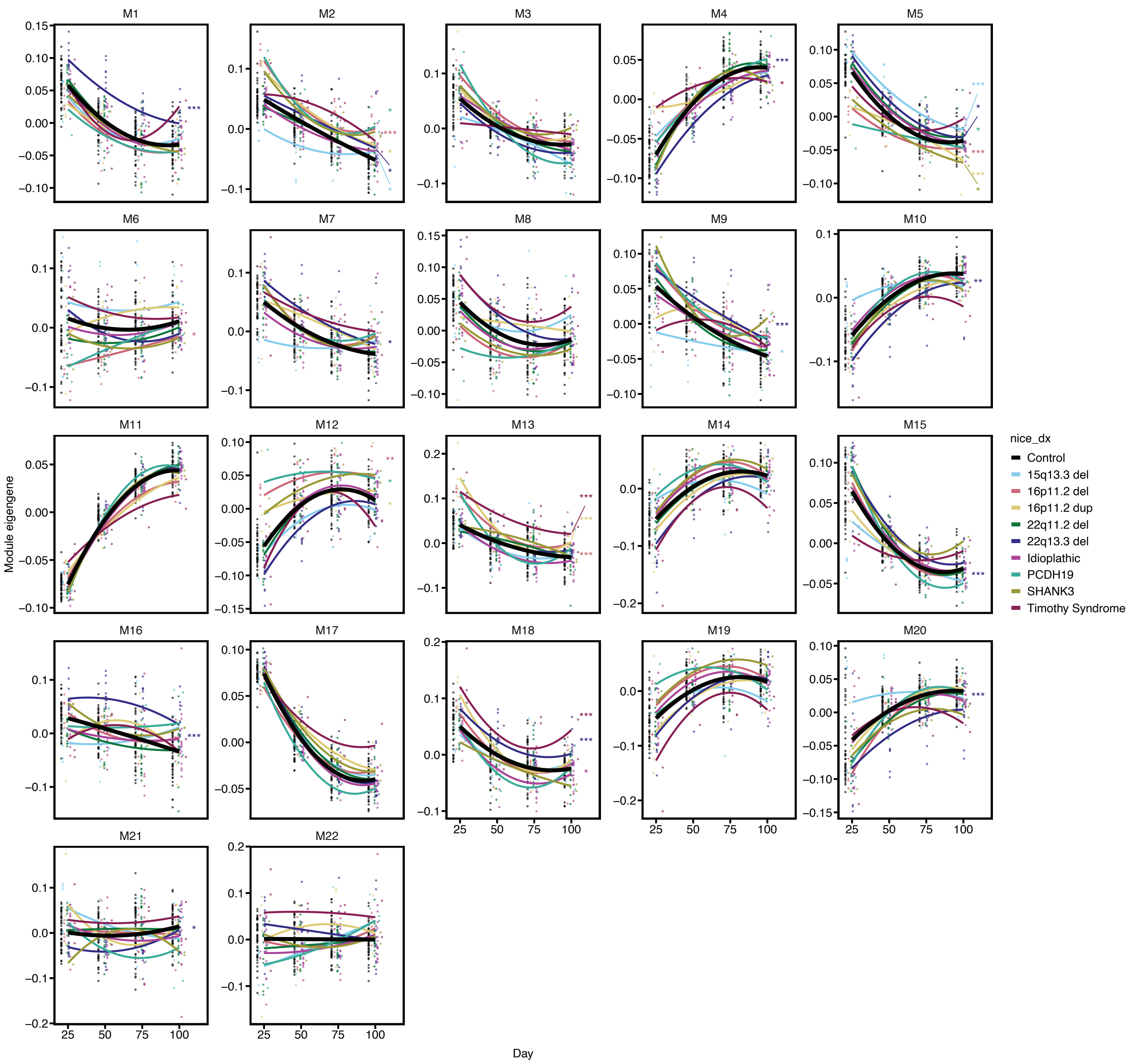


**b**

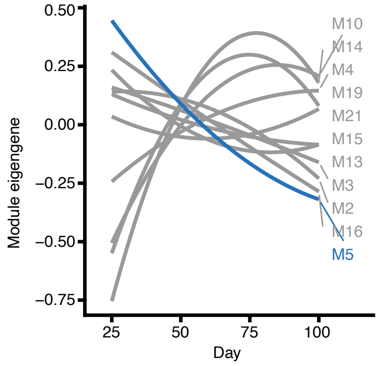

Extended data figure 14

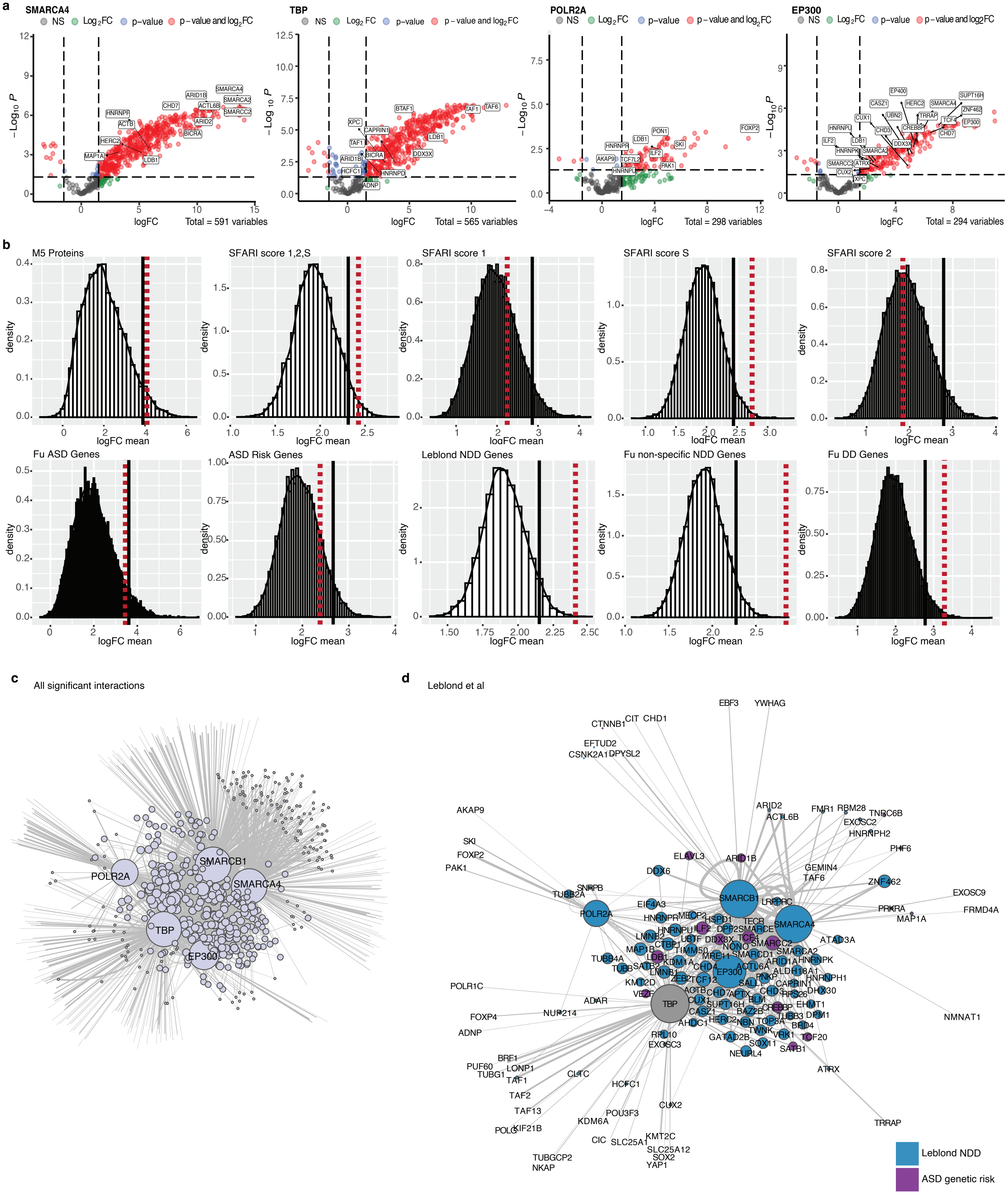

Extended Data Figure 15

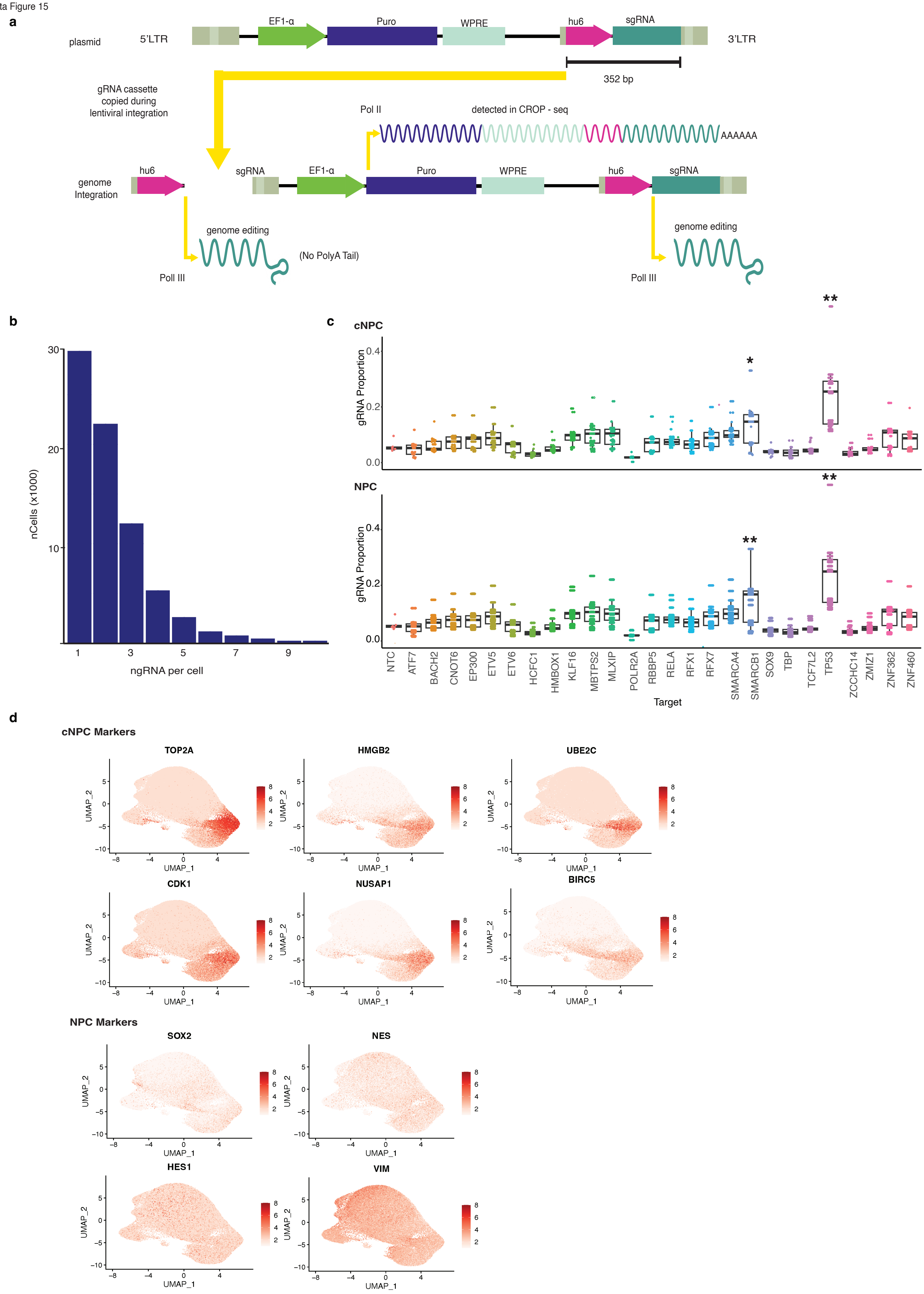

Extended Data Figure 16

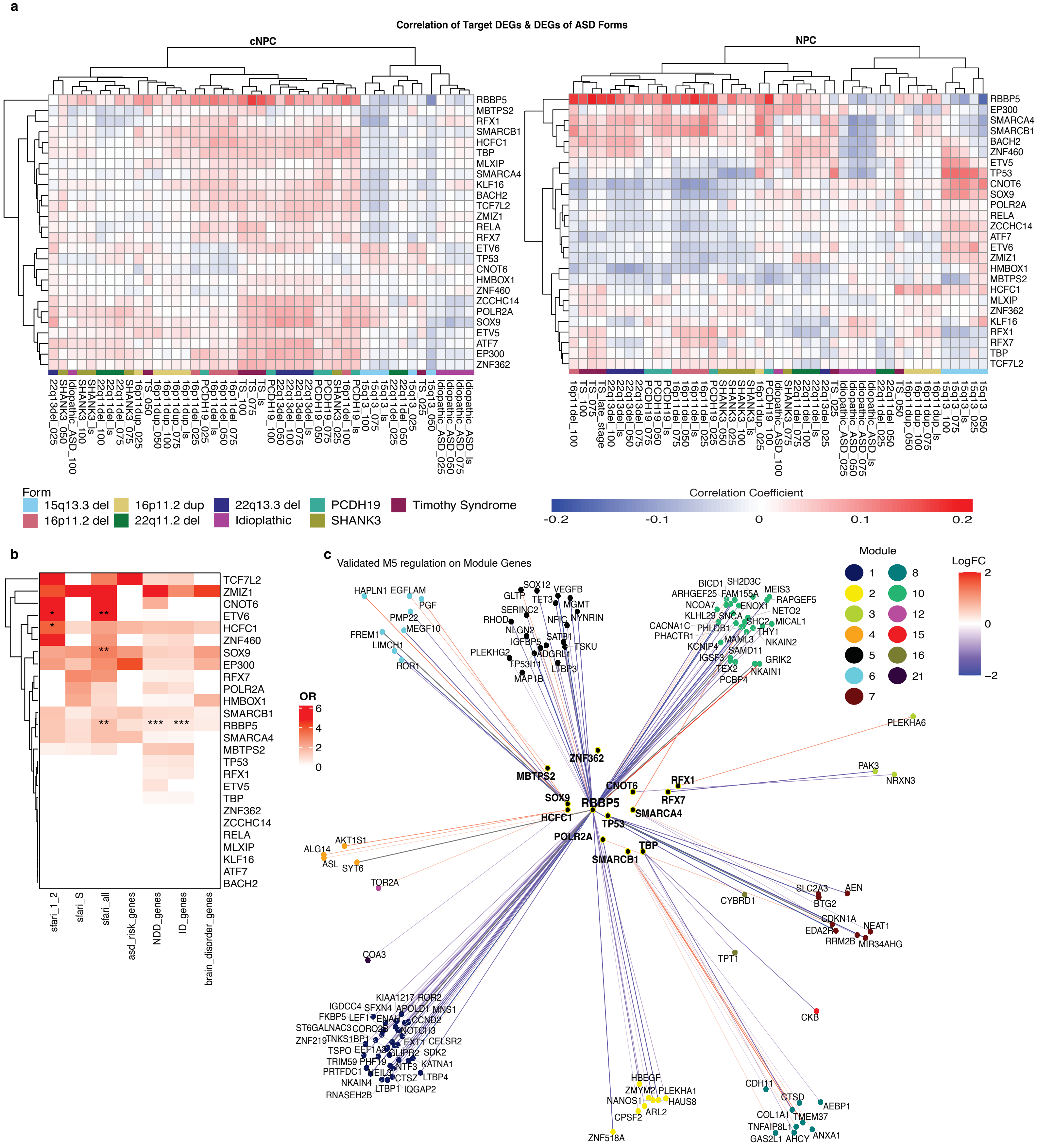

Extended Data Figure 17

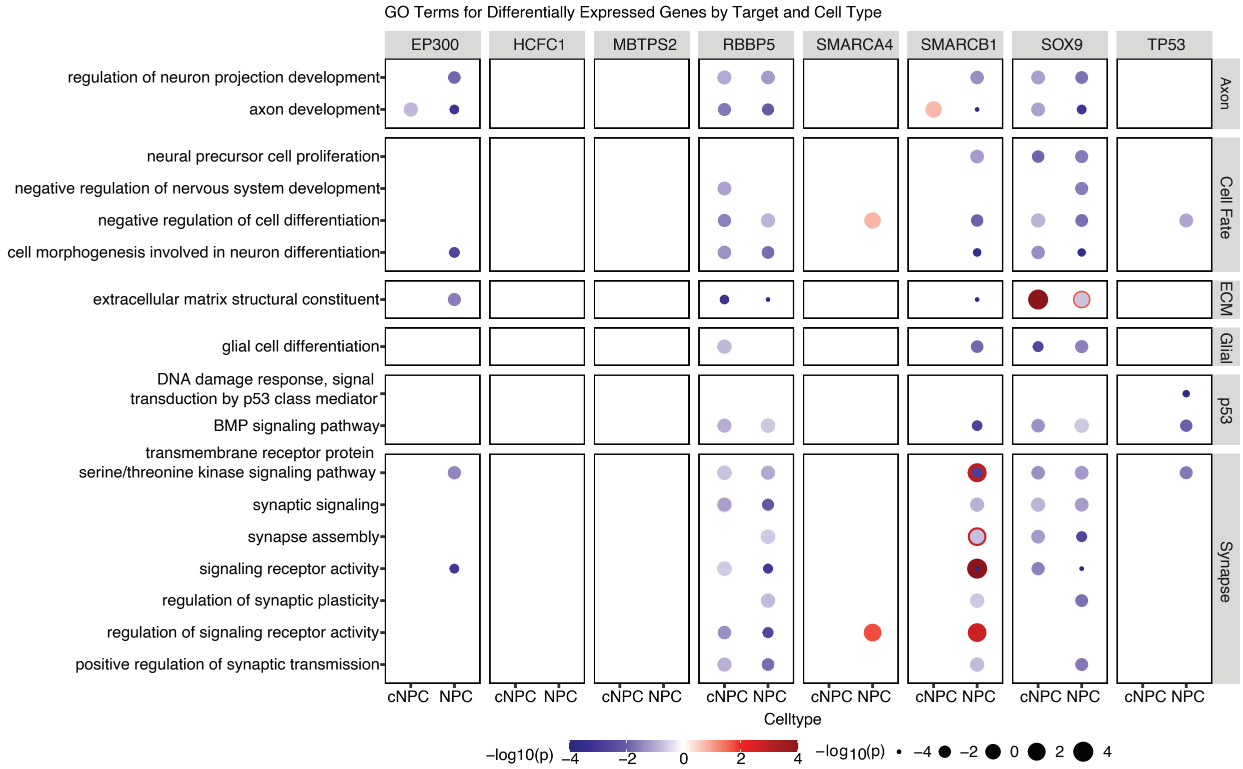
